# The Paf1 complex broadly impacts the transcriptome of *Saccharomyces cerevisiae*

**DOI:** 10.1101/567495

**Authors:** Mitchell A. Ellison, Alex R. Lederer, Marcie H. Warner, Travis Mavrich, Elizabeth A. Raupach, Lawrence E. Heisler, Corey Nislow, Miler T. Lee, Karen M. Arndt

## Abstract

The Polymerase Associated Factor 1 complex (Paf1C) is a multifunctional regulator of eukaryotic gene expression important for the coordination of transcription with chromatin modification and post-transcriptional processes. In this study, we investigated the extent to which the functions of Paf1C combine to regulate the *Saccharomyces cerevisiae* transcriptome. While previous studies focused on the roles of Paf1C in controlling mRNA levels, here we took advantage of a genetic background that enriches for unstable transcripts and demonstrate that deletion of *PAF1* affects all classes of Pol II transcripts including multiple classes of noncoding RNAs. By conducting a *de novo* differential expression analysis independent of gene annotations, we found that Paf1 positively and negatively regulates antisense transcription at multiple loci. Comparisons with nascent transcript data revealed that many, but not all, changes in RNA levels detected by our analysis are due to changes in transcription instead of post-transcriptional events. To investigate the mechanisms by which Paf1 regulates protein-coding genes, we focused on genes involved in iron and phosphate homeostasis, which were differentially affected by *PAF1* deletion. Our results indicate that Paf1 stimulates phosphate gene expression through a mechanism that is independent of any individual Paf1C-dependent histone modification. In contrast, the inhibition of iron gene expression by Paf1 correlates with a defect in H3 K36 tri-methylation. Finally, we showed that one iron regulon gene, *FET4*, is coordinately controlled by Paf1 and transcription of upstream noncoding DNA. Together these data identify roles for Paf1C in controlling both coding and noncoding regions of the yeast genome.

## INTRODUCTION

In the context of chromatin, accurate and controlled transcription by RNA polymerase II requires the functions of many regulatory factors. One highly conserved regulatory factor is Paf1C, which in yeast is composed of Paf1, Ctr9, Leo1, Rtf1, and Cdc73 (Jaehning 2010; Crisucci and Arndt 2011; Tomson and Arndt 2013). Paf1C associates with Pol II during transcription elongation and regulates both transcriptional and post-transcriptional processes, including the co-transcriptional deposition of histone modifications and the nuclear export of RNAs (Tomson and Arndt 2013; Fischl *et al.* 2017; Van Oss *et al.* 2017). Histone modifications dependent on Paf1C include H3 lysine 4 di- and tri-methylation (H3 K4me2/3), H3 K79me2/3, and H2B K123 mono-ubiquitylation (ub) in *S. cerevisiae* (H2B K120 in humans). Paf1C facilitates the deposition of H2B K123ub via its Rtf1 subunit (Piro *et al.* 2012; Van Oss *et al.* 2016), which directly interacts with the ubiquitin-conjugating enzyme Rad6 through its histone modification domain (Van Oss *et al.* 2016). H2BK123ub is a prerequisite to H3 K4me2/3 and H3 K79me2/3, modifications catalyzed by the histone methyltransferases Set1 and Dot1, respectively (Dover *et al.* 2002; Sun and Allis 2002). Paf1C also promotes the deposition of H3 K36me3 by the Set2 histone methyltransferase (Chu *et al.* 2007). Consistent with the binding of Paf1C to the Pol II elongation machinery (Qiu *et al.* 2006, 2012; Amrich *et al.* 2012; Wier *et al.* 2013; Xu *et al.* 2017; Vos *et al.* 2018), the histone modifications dependent on Paf1C are found at regions of active transcription (Smolle and Workman 2013).

The absence of specific histone modifications in Paf1C mutants is associated with transcriptional defects. These defects include the transcriptional read-through of terminators found at the 3’ ends of Pol II-transcribed small nucleolar RNA (snoRNA) genes (Sheldon *et al.* 2005; Terzi *et al.* 2011; Tomson *et al.* 2011, 2013). In addition to promoting a histone modification state that facilitates transcription termination, Paf1C physically associates with proteins implicated in transcription termination and RNA 3’-end formation (Nordick *et al.* 2008; Rozenblatt-Rosen *et al.* 2009). The importance of Paf1C in regulating snoRNA termination supports a functional interaction with the Nrd1-Nab3-Sen1 (NNS) transcription termination pathway (Arndt and Reines 2014; Porrua and Libri 2015), which is responsible for the termination of snoRNAs and other noncoding transcripts including cryptic unstable transcripts (CUTs) in yeast (Schulz *et al.* 2013). Many of these same noncoding transcripts are rapidly degraded by the nuclear exosome through a process mediated by the Trf4/Trf5-Air1/Air2-Mtr4 polyadenylation complex (TRAMP) (Schmid and Jensen 2008). For example, loss of Trf4, the polyA polymerase subunit of the TRAMP complex that adds short polyA tails to transcripts destined for degradation or processing by the nuclear exosome, has been shown to stabilize CUTs and snoRNAs in *S. cerevisiae* (LaCava *et al.* 2005; Vanácová *et al.* 2005; Wyers *et al.* 2005; Thiebaut *et al.* 2006; Fallis 2009; Xu *et al.* 2009).

Despite growing understanding of the molecular functions of Paf1C, few studies have probed how these functions lead to a transcriptional outcome. Moreover, little is known about the roles of Paf1C in controlling noncoding transcription. To begin to address these questions, we sought to comprehensively investigate the importance of Paf1C in modulating the *S. cerevisiae* transcriptome, taking advantage of a genetic background that allows enhanced detection of unstable transcripts. To this end, we used strand-specific whole-genome tiling arrays to measure steady state RNA levels in *PAF1* and *paf1Δ* strains that contain or lack the TRAMP subunit gene *TRF4*. We found that deletion of *PAF1* affects all classes of Pol II transcripts including both stable and unstable noncoding RNAs and antisense transcripts. Comparisons with published NET-seq experiments, which detect Pol II-engaged, nascent transcripts (Harlen and Churchman 2017), indicate that most, but not all, changes in steady state transcript abundance in the *paf1Δ* background can be attributed to altered transcription. Analysis of subsets of protein-coding genes suggests that Paf1 represses the transcription of some genes through facilitating H3 K36me3 and stimulates the transcription of other genes independently of any single Paf1C-dependent histone modification. Finally, we report a regulatory mechanism governing the *FET4* locus, which incorporates both CUT transcription and Paf1. Together these data support a role for Paf1C in multiple regulatory mechanisms that collectively and broadly impact the Pol II transcriptome.

## MATERIALS AND METHODS

### Yeast strains and culturing methods

All *S. cerevisiae* strains used in this study are listed in Table 1 and are isogenic to the FY2 strain, which is a *GAL2*^*+*^ derivative of S288C (Winston *et al.* 1995). Deletion of specific loci was achieved by one step gene disruption (Lundblad *et al.* 2001) and confirmed by PCR. Genetic crosses were conducted as described (Rose *et al.* 1991). Cells were grown to log phase at 30°C in rich media (YPD) supplemented with 400 µM tryptophan and harvested by centrifugation. Cell pellets were washed once with sterile water, flash frozen, and stored at −80°C prior to RNA isolation for RT-qPCR and Northern blotting experiments.

**Table 1.**
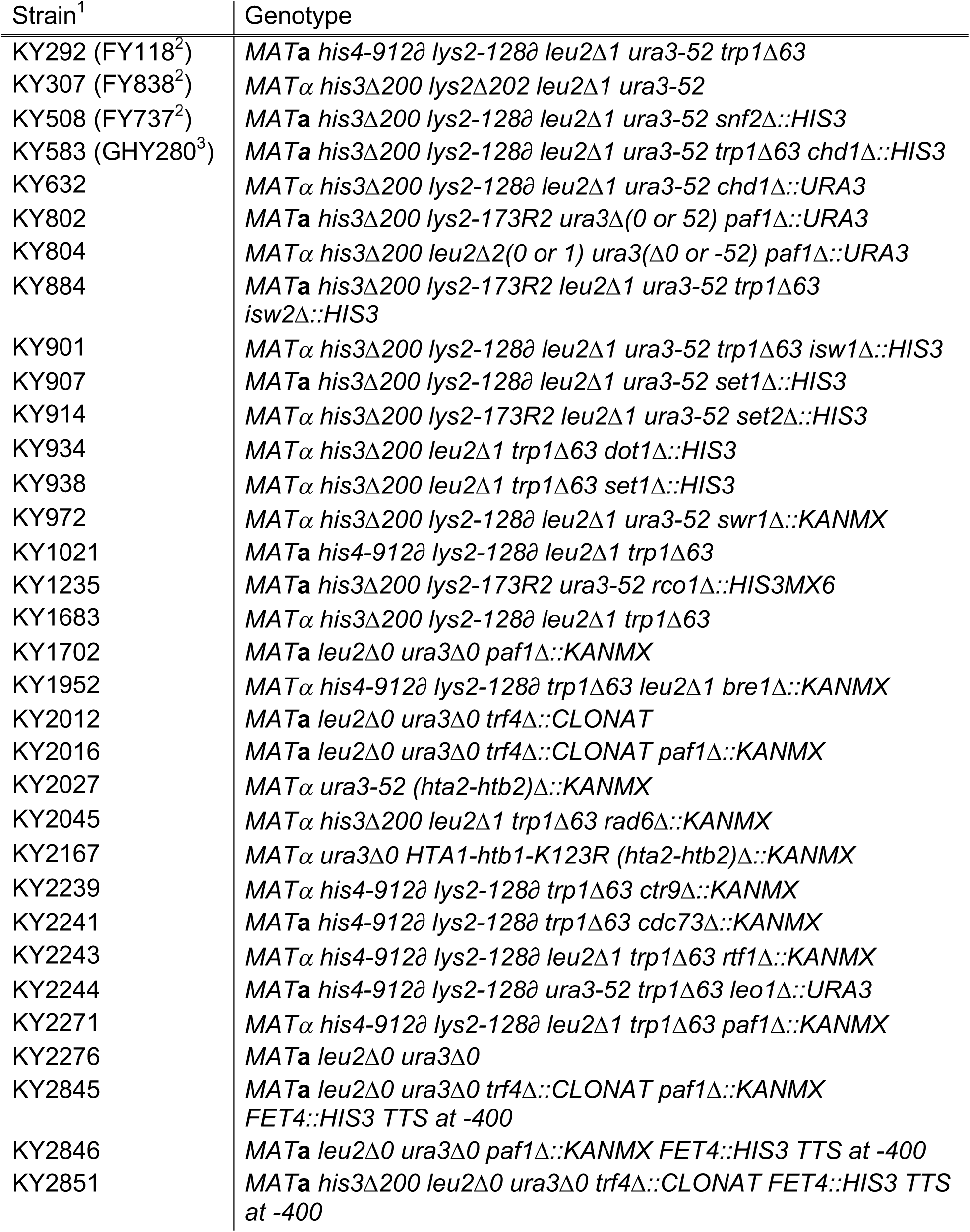
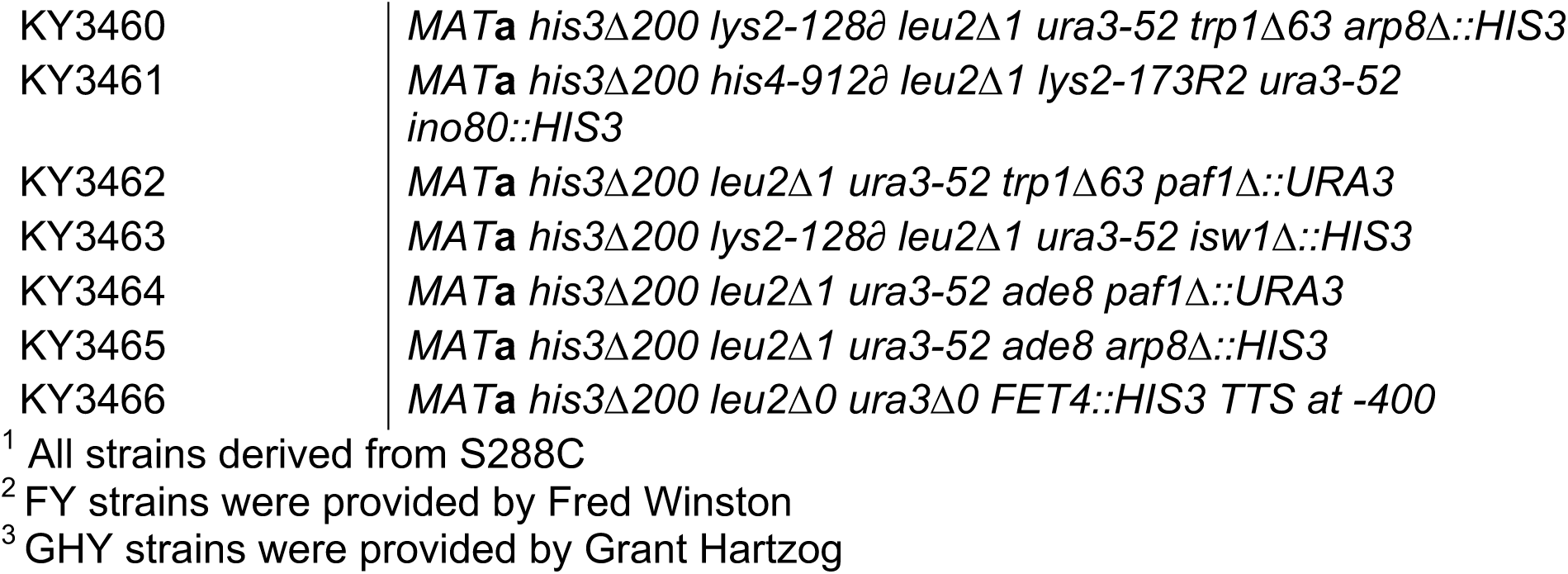
Yeast strains used in this study

### RNA isolation

RNA was extracted by the hot phenol extraction method (Collart and Oliviero 1993). Briefly, frozen cells were suspended in 400 μL of TES extraction buffer (10 mM EDTA, 10 mM Tris-Cl pH 7.5, 0.5% SDS) and 400 μL of acid phenol, followed by incubation at 65°C for 1hr. The aqueous phase was collected and re-extracted using acid phenol and then chloroform. Extracted RNA was combined with 40 μL of 3 M sodium acetate and 1 mL of 100% ethanol, mixed, and placed at −80°C for at least 1 hr. Precipitated RNA was collected by centrifugation and suspended in RNase-free water before quantification and quality check by agarose gel electrophoresis.

### Northern blot analysis

For Northern blot analyses, 10μg-20µg of total RNA were separated on a gel containing 2% agarose, 6.5% formaldehyde, and 1X MOPS for 500 volt hr and then transferred to a Nytran supercharge nylon transfer membrane (Schleicher & Schuell BioScience, #10416296, Dassel, Germany) prior to hybridization with radiolabeled DNA probes. DNA probes were generated by PCR corresponding to the following genomic regions relative to the +1 nucleotide of the annotated coding sequence for the *FET4* gene: *CUT 793/794* (−479 to −114) and *FET4* (+261 to +651). Detection of *SCR1* (−181 to +284) served as a loading control. Oligonucleotides used to generate these probes are listed in Table S1. Probes were made using [*α*^32^P]-dATP (single labeling) or [*α*^32^P]-dATP and [*α*^32^P]-dTTP (double labeling). Signals were quantified using a Typhoon FLA 7000 phosphorimager (GE, Boston, MA) and normalized to the *SCR1* internal loading control using ImageJ software.

### Reverse transcription quantitative polymerase chain reaction (RT-qPCR)

A total of 10 µg of RNA from each sample to be used in RT-qPCR was treated with TURBO DNase (Ambion, AM1907, Thermo Fisher, Waltham, MA) following manufacturer’s instructions. To ensure that there was no DNA contamination after the DNase treatment, 1 μL of DNase-treated RNA was subjected to 40 cycles of PCR and analyzed by agarose gel electrophoresis. All samples used in RT-qPCR showed no PCR product after 40 cycles. Reverse transcription reactions were performed on 1 µg of DNase-treated RNA using the RETROScript Reverse Transcription Kit (Ambion, AM1710, Thermo Fisher, Waltham, MA) following the manufacturer’s instructions.

RT-qPCR experiments were performed in technical duplicate and all strains were tested in at least biological triplicate. Reactions were prepared in a volume of 20 µL using Maxima SYBR Green/ROX qPCR Master Mix (2X) (Thermo Fisher # K0221, Waltham, MA) following the manufacturer’s instructions. Each 20 μL reaction was then divided into two 10 μL reactions, which were analyzed on a StepOne™ Real-Time PCR System (Thermo Fisher, Waltham, MA) beginning with a hold at 95°C for 10 min followed by 40 cycles of 95°C for 15 sec and 58°C for 1 min and finally terminating with the generation of a melt curve. Efficiencies were determined for all primer sets by measuring Ct values across a series of six ten-fold dilutions starting with 250 ng/μL and ending with 2.5 pg/μL. RT-qPCR data were analyzed using the mathematical formula developed by (Pfaffl 2001) and normalized to *SCR1* levels. RT-qPCR primers and their efficiencies are listed in Table S1.

### Affymetrix tiling array analysis

All RNA samples used in tiling array analysis were prepared using established methods (Juneau *et al.* 2007; Perocchi *et al.* 2007) and quality was assessed by agarose gel electrophoresis. Briefly, 100 µg RNA were treated with DNase (Fermentas #EN0521, Waltham, MA) and purified using an RNeasy kit (Qiagen #74104, Hilden Germany). Then 25 µg of RNA were reverse transcribed into cDNA using 1.8 kU SuperScript II RT (Invitrogen #18064-014), 12.5 ng/µL random hexamers, and 12.5 ng/µL oligo(dT) at 42°C for 2 h in the presence of 6 ng/µL actinomycin D (Sigma, #A1410-2MG) to prevent antisense artifacts (Perocchi *et al.* 2007). RNA was removed using NaOH hydrolysis and cDNA was purified using the buffers from the QIAquick Nucleotide Removal Kit and the columns from the MinElute Reaction Cleanup Kit (Qiagen). Finally, cDNA was fragmented and labeled with a 3’ biotin tag before quantifying gene expression using Affymetrix custom tiling arrays (A-AFFY-116 - Affymetrix Custom Array - *S. cerevisiae* Tiling Steinmetz, GEO Platform ID: GPL4563) (David *et al.* 2006; Huber *et al.* 2006).

### Generation of annotation files for the tilingArray R package

Following the guidelines provided with the *davidTiling* Bioconductor package (David *et al.* 2006), tiling array probes were mapped to the *S. cerevisiae* genome (S288C version = R64-2-1) (Cherry *et al.* 2012; Engel *et al.* 2014). Briefly, the probe FASTA file was extracted from the array design file and used as input for MUMmer3.23 (Kurtz *et al.* 2004), along with the chromosome FASTA files for the S288C genome. Both the output of MUMmer3.23 and the S288C genome annotations were read into R (Team 2016), and a slightly modified version of the *makeProbeAnno.R* script was used to generate an up-to-date probe annotation file for use with the *tilingArray* package (Huber *et al.* 2006). All R packages used in this study can be found at www.bioconductor.org (Gentleman *et al.* 2004) and R scripts can be found at https://github.com/mae92/Paf1C-Transcriptome-Analysis-Code/tree/master/R_Code.

### Variance stabilizing normalization

CEL files for 12 tiling arrays were read into R as a single expression set using the *readCel2eSet()* function of the *tilingArray* package. The probe intensities for all 12 arrays were log2 transformed and normalized (variance stabilized normalization R code available at: https://github.com/mae92/Paf1C-Transcriptome-Analysis-Code/tree/master/R_Code) to minimize batch effects using the *vsn* package (Huber *et al.* 2002).

### Mapping probe intensities to probe positions across the *S. cerevisiae* genome

The expression set containing the normalized log2 transformed probe intensities was used as input for the *segChrom()* function of the *tilingArray* package, and the locations of probes across the genome were extracted for use in downstream analysis using basic R commands (R code can be found at: https://github.com/mae92/Paf1C-Transcriptome-Analysis-Code/tree/master/R_Code). Probe locations were averaged for triplicate samples and these averaged values were used to generate BedGraph files, which were converted into bigWig files for visualization in the Integrative Genomics Viewer (IGV) from the Broad Institute (Thorvaldsdottir *et al.* 2013).

### Annotation-guided differential expression analysis

Normalized log2 transformed probe intensity values were extracted from the *tilingArray* output. Using a custom file (available at: https://github.com/mae92/Paf1C-Transcriptome-Analysis-Code/blob/master/Transcript_Annotations/combined.fix.csv) containing transcript annotations (listed in Table S2) from the Saccharomyces Genome Database (SGD) and recent studies of novel noncoding RNA transcripts (Cherry *et al.* 1998; Xu *et al.* 2009; Yassour *et al.* 2010; van Dijk *et al.* 2011; Schulz *et al.* 2013; Venkatesh *et al.* 2016), we calculated the average log2 intensity values for probes spanning each annotation. This process was carried out using an in-house Python script (available at: https://github.com/mae92/Paf1C-Transcriptome-Analysis-Code/tree/master/Python_Code) that calculates the average intensity of all probes occupying a given annotation found in the annotation file. Average log2 intensity values for all transcripts in each replicate and strain background were loaded into the *limma* package where a linear model was used to determine statistical significance. Log2 fold change and *p* values for all transcripts tested were extracted from the output for each strain comparison. These *p* values were adjusted using Benjamini and Hochberg’s false discovery rate (FDR) (Benjamini and Hochberg 1995) method using *top.table* command in *limma* (R code for *limma* analysis can be found here: https://github.com/mae92/Paf1C-Transcriptome-Analysis-Code/tree/master/R_Code) (Ritchie *et al.* 2015). Significantly differentially expressed genes (adjusted *p* value < 0.05) that were present in both comparisons (*paf1Δ* vs WT and *paf1Δ trf4Δ* vs *trf4Δ)* were loaded into SGD’s YeastMine database (Balakrishnan *et al.* 2012), where additional annotation and gene ontology information could be extracted. Gene ontology results are shown in Table 2. Plots of differential expression data were produced in R.

**Table 2.**
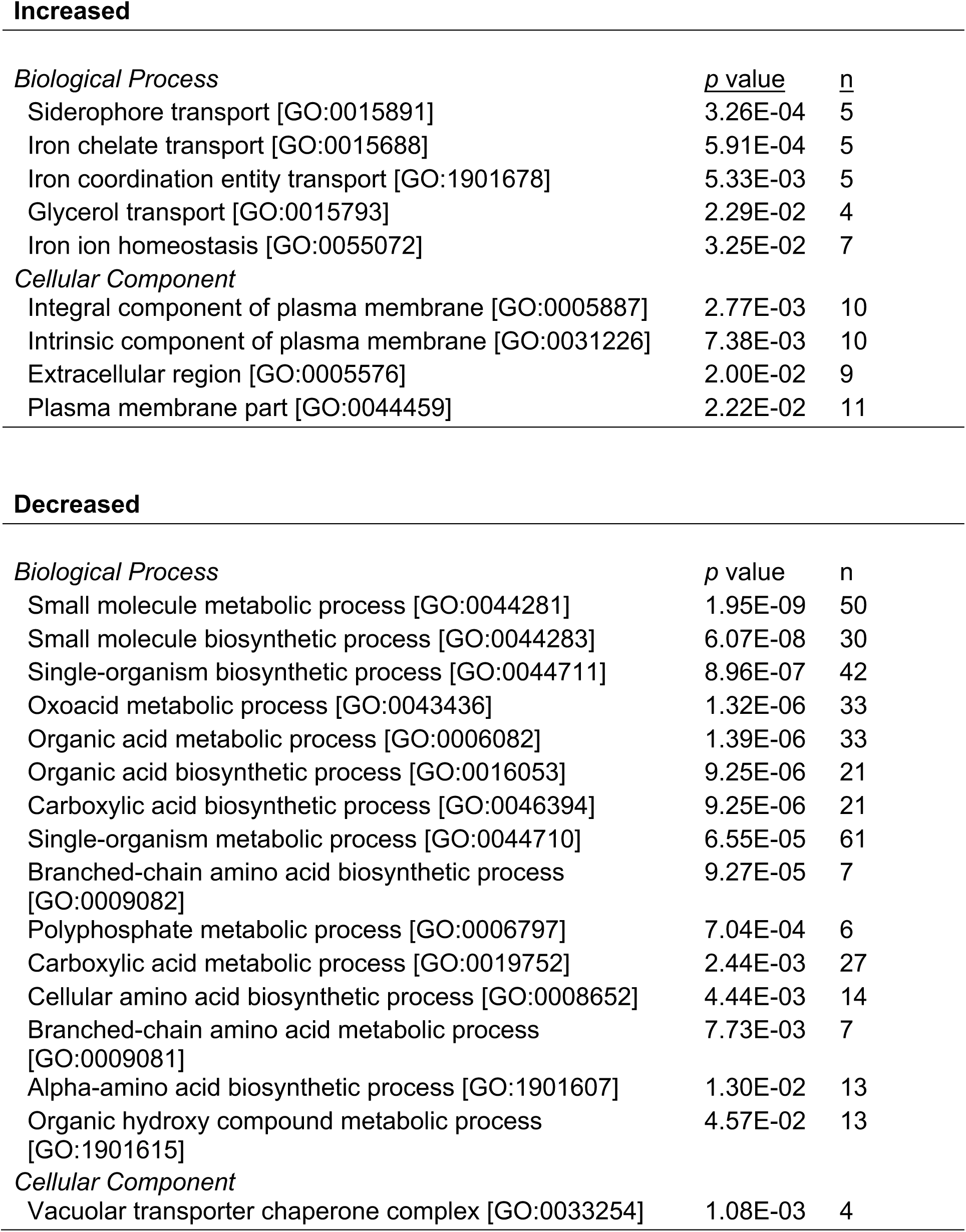
Gene ontology results for genes that showed increased or decreased expression (1.5-fold) in both *paf1Δ* to WT and *paf1Δ trf4Δ* to *trf4Δ* comparisons

### *De novo* differential expression analysis

BedGraph files were created containing normalized log2 probe intensity values (averaged for the three biological replicate arrays) mapped to the yeast genome. Differentially expressed transcripts identified by the *de novo* differential expression analysis were defined by a six-step process (Figure S1). 1) Average log2(probe intensity) was calculated for three biological replicates. 2) The data were smoothed by averaging across a sliding window of 20 tiling array probes (roughly 160bp). 3) The log2 fold change (experimental vs. control) was calculated across the entire genome. 4) All regions with an absolute fold change greater than one (log2 fold change of 0) were identified. 5) Regions of the genome with an absolute change of 1.5-fold (log2 fold change of 0.58) were identified and any of these regions less then 80 bp long (a length comparable to the shortest snoRNA) were excluded. 6) The two lists of regions from steps 4 and 5 were intersected to yield a list of extended differentially expressed regions where some portion of the transcript had an absolute fold change of 1.5-fold or greater. The transcripts defined by this method were then treated as their own list of transcript annotations for use in the comparisons described herein. This was done using a combination of AWK (Aho *et al.* 1979) and the BedTools suite (Quinlan and Hall 2010; Quinlan 2014). The shell script used to define differentially expressed transcripts can be found at https://github.com/mae92/Paf1C-Transcriptome-Analysis-Code/tree/master/Shell_Code and R code used to generate the input BedGraph files can be found at https://github.com/mae92/Paf1C-Transcriptome-Analysis-Code/tree/master/R_Code.

### Analysis of published datasets

Next generation sequencing datasets from previous studies (Churchman and Weissman 2011; Van Oss *et al.* 2016; Harlen and Churchman 2017) were obtained in BedGraph format directly from the authors or FASTQ format from NCBI SRA database and converted into BigWig format for use with DeepTools (Ramírez *et al.* 2014, 2016). Files received in BedGraph format were converted using the University of California Santa Cruz (UCSC) Genome Browser (Kent *et al.* 1976) utility *BedGraphToBigWig.* Files downloaded from the SRA database in FASTQ format were mapped to the *S. cerevisiae* genome (S288C version = R64-2-1) (Cherry *et al.* 2012; Engel *et al.* 2014) using HISAT2 (Kim *et al.* 2015) and converted to BAM format using Samtools (Li *et al.* 2009). BAM files were converted to Wig format using the bam2wig utility (found at https://github.com/MikeAxtell/bam2wig) and converted to BigWig format using the UCSC utility *WigToBigWig.* Heatmaps were plotted using *computeMatrix* and *plotHeatmap* tools in the deepTools package by summing the tag counts using 50bp bins.

### Statistical Analysis

At least three biological replicates were performed for every assay shown in this manuscript including tiling arrays. Each biological replicate is a pure yeast culture derived from a single colony initiated from a single cell of a given strain. Tiling array data analyzed using the *limma* package were subjected to the standard *limma* workflow which utilizes linear modeling and an empirical Bayes method to determine differentially expressed genes from as little as three biological replicates. The *limma p* values were adjusted for multiple comparisons using Benjamini and Hochberg’s false discovery rate (FDR) (Benjamini and Hochberg 1995). All RT-qPCR and Northern blot *p* values were generated using an unpaired, two-sided, students t-test assuming equal variance carried out between the mutant strain and the wild type strain.

### Data Availability

Strains are available upon request. Tiling array data (raw CEL files, BedGraph files, annotation-guided differential expression results, and files containing annotations for differentially expressed transcripts defined by our *de novo* analysis in BED6 file format) have been deposited in the Gene Expression Omnibus database under accession number GSE122704. Code used for analysis of tiling array data has been uploaded to the following GitHub repository (https://github.com/mae92/Paf1C-Transcriptome-Analysis-Code). Tables S1-S6 and Figures S1-S5 are available via FigShare.

## RESULTS

### Deletion of *PAF1* affects coding and noncoding transcripts genome-wide

To investigate the impact of Paf1C on the *S. cerevisiae* transcriptome, we used high-resolution whole-genome tiling arrays to measure steady state RNA levels in *S. cerevisiae* strains deleted for the *PAF1* gene, which encodes a core member of Paf1C important for complex integrity (Mueller *et al.* 2004; Deng *et al.* 2018). Additionally, to assess the Paf1-dependency of unstable noncoding RNAs (ncRNAs) in these experiments, we deleted *TRF4* in both *PAF1* and *paf1Δ* strains. When compared to a *trf4Δ* strain, the *paf1Δ trf4Δ* double mutant revealed wide-ranging effects on all Pol II transcript classes examined: mRNAs, small nuclear RNAs (snRNAs), snoRNAs, CUTs, stable unannotated transcripts (SUTs; Xu *et al.* 2009), Xrn1-dependent unstable transcripts (XUTs; van Dijk *et al.* 2011), Nrd1-unterminated transcripts (NUTs; Schulz *et al.* 2013), and Set2-repressed antisense transcripts (SRATs; Venkatesh *et al.* 2016) (Figure 1A-G; Table S3). In general, levels of snRNAs, snoRNAs, and SRATs increased in the *paf1Δ trf4Δ* double mutant relative to the *trf4Δ* single mutant (Figures 1D and G; Table S3) indicating that, in wild type cells, Paf1 suppresses their transcription or destabilizes the transcripts. In the case of SRATs, increased transcript levels are consistent with a requirement for Paf1 in facilitating H3 K36me3, a modification that negatively regulates transcription (Churchman and Weissman 2011; Kim *et al.* 2016; Venkatesh *et al.* 2016) by activating the Rpd3S histone deacetylase complex and by inhibiting histone exchange (Carrozza *et al.* 2005; Joshi and Struhl 2005; Keogh *et al.* 2005; Govind *et al.* 2010; Venkatesh *et al.* 2012). Levels of many CUTs, SUTs, XUTs and NUTs decreased upon deletion of *PAF1* (Figures 1B, C, E and F; Table S3). For NUTs and CUTs, these changes in transcript abundance suggest that Paf1 impacts NNS-dependent termination beyond the snoRNA genes. At protein-coding genes, Paf1 positively and negatively affects mRNA levels in a locus-specific manner (Figure 1A; Table S3), in agreement with previous studies (Shi *et al.* 1996; Porter *et al.* 2005; Cao *et al.* 2015; Chen *et al.* 2015; Yu *et al.* 2015; Yang *et al.* 2016; Fischl *et al.* 2017).

**Figure 1.**
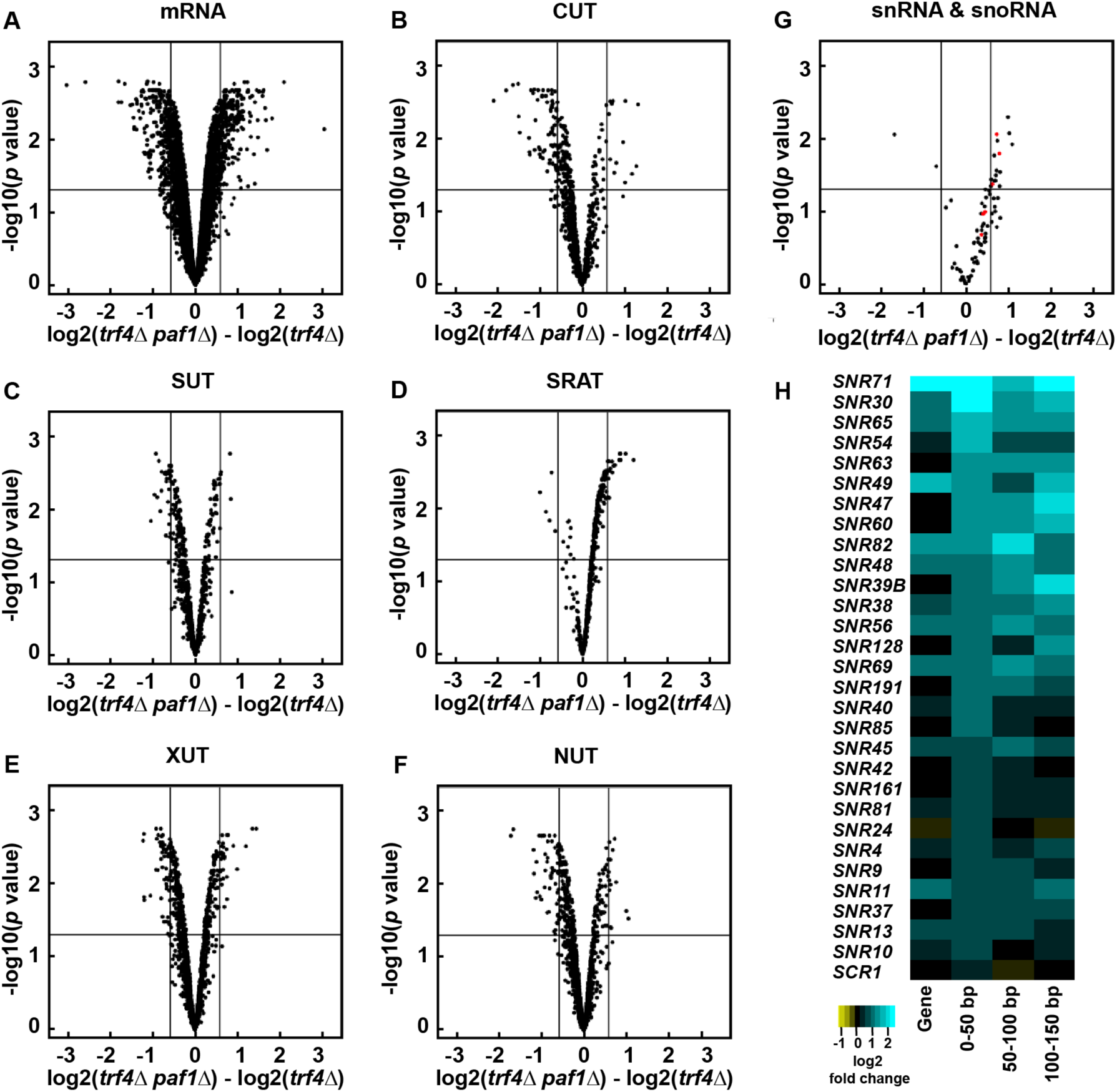
Deletion of *PAF1* affects all Pol II transcript classes. (A-G) Volcano plots graphing statistical significance (y-axis) against expression change (x-axis) between *paf1Δ trf4Δ* and *trf4Δ* strains (KY2016 and KY2012, respectively) for the indicated Pol II transcript classes. In panel G, snRNAs and snoRNAs are shown in red and black, respectively. Each point represents an individual transcript. Tiling array probe intensities were averaged over annotated regions using a custom Python script and an average log2 fold change and *p* value were calculated using the *limma* R package. The horizontal line indicates an FDR adjusted *p* value of 0.05 and the vertical lines indicate a 1.5-fold change in expression (log2 fold change of 0.58). Counts and percentages of differentially expressed transcripts shown here are listed in Table S3. (H) Heatmap of log2 fold change in expression between *paf1Δ* and WT strains (KY1702 and KY2276, respectively) for the 29 most affected snoRNA genes. The snoRNA gene bodies and regions 0-50 bp, 50-100 bp, and 100-150 bp downstream of their annotated 3’ ends are plotted and sorted by the 0-50 bp region.

For many snoRNA genes, we detected an increase in RNA levels downstream of the annotated gene in the *paf1Δ* strain relative to wild type, consistent with previous studies showing Paf1 is required for efficient snoRNA termination (Sheldon *et al.* 2005; Tomson *et al.* 2013). The log2 fold change values calculated for any particular snoRNA gene and its downstream region do not always agree. In many cases, RNA levels mapping to the gene body do not change expression even when downstream changes are observed, suggesting that read-through transcription is occurring at these loci (Figure 1H, compare gene and 0-50bp heat map columns).

### *De novo* differential expression analysis reveals effects of Paf1 on antisense transcripts

As an independent analysis and to facilitate detection of unannotated transcripts, we performed a *de novo* differential expression analysis of our tiling array data. Here, we relied on the data to reveal the boundaries of differentially expressed transcripts instead of using predetermined annotations. Strains lacking *PAF1* were compared to control strains on a probe-by-probe basis. Genomic regions with a 1.5-fold or greater difference in expression between *paf1Δ* and *PAF1* strains were selected as differentially expressed and extended until the expression difference was no longer observed (see Figure S1 and Materials and Methods for a detailed description of this analysis).

Confirming the accuracy of the *de novo* analysis, we found that nearly all the mRNAs identified as differentially expressed in the *paf1Δ trf4Δ* strain by our annotation-guided analysis (585 mRNAs;1.5-fold or greater expression change relative to *trf4Δ* strain) were also detected by the *de novo* analysis (Figure S2A). We note that, compared to the annotation-guided analysis, a larger number of differentially expressed transcripts that overlap mRNAs on the sense strand were detected in the *de novo* analysis. This observation is not due to technical differences, but rather a consequence of multiple *de novo* transcripts overlapping with a single annotated mRNA and the increased sensitivity of the analysis. Additionally, we found that the length distribution of all transcripts identified in the *de novo* analysis was similar to that reported in SGD for mRNAs (Figure S2B), confirming that we were not calling exceedingly long or short transcripts. Further, separation of the *de novo* analysis data by transcript class revealed effects of *paf1Δ* on noncoding and coding RNAs similar to those observed in the annotation-guided analysis (compare Figure 1A-1G to Figure 2A; Table S4). In the *paf1Δ trf4Δ* strain, for example, SRATs and snoRNAs were predominantly up-regulated and other noncoding RNAs were predominantly down-regulated. When viewed as a whole, the *de novo* analysis detected far more differentially expressed transcripts in the *paf1Δ trf4Δ* strain (relative to the *trf4Δ* control strain) than in the *paf1Δ* strain (relative to the *PAF1* control strain) (Figure S2C). Therefore, a functional TRAMP complex, which promotes processing and degradation of unstable transcripts, obscures many of the transcriptional effects of deleting *PAF1*.

**Figure 2.**
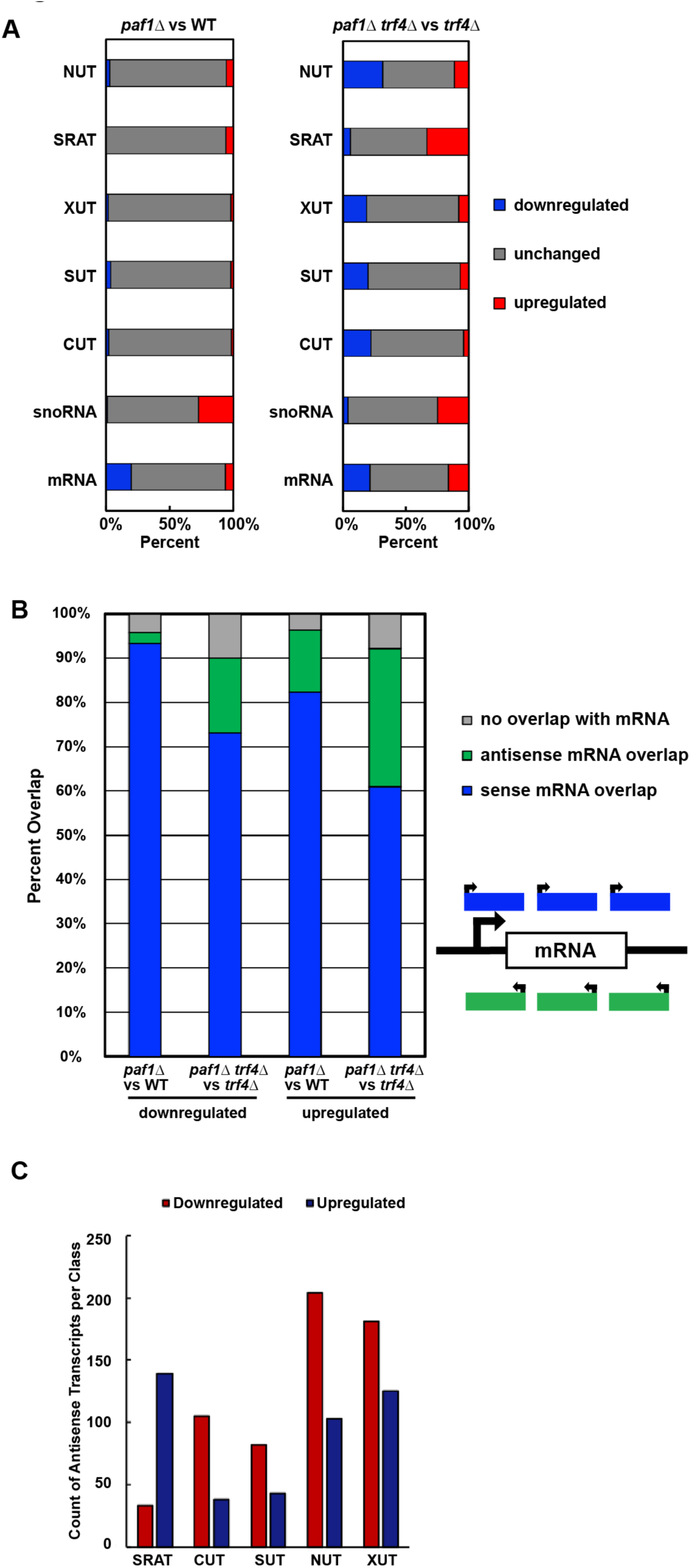
Paf1 positively and negatively regulates antisense transcription. (A) Horizontally stacked bar graphs showing the percentage of each transcript class (listed in Table S2) found to overlap with a differentially expressed transcript identified in *paf1Δ* or *paf1Δ trf4Δ* strains by *de novo* analysis (counts and percentages listed in Table S4). (B) Vertically stacked bar graph plotting percentage of transcripts, identified in the *de novo* analysis, that overlap with mRNA coding regions on the sense or antisense strand. These data are also presented in Table S5. (C) Bar graph summarizing the overlap between differentially expressed antisense transcripts detected by the *de novo* analysis and previously annotated noncoding RNAs (see sums in Table S6 for counts).

Unstable ncRNAs are often found near mRNA loci in tandem or antisense orientations (Neil *et al.* 2009; Xu *et al.* 2009; van Dijk *et al.* 2011; Schulz *et al.* 2013; Castelnuovo *et al.* 2014). Murray *et al.* (2015) demonstrated that regions of high antisense transcription are deficient in H2BK123ub, H3K4me3, H3K79me3 and H3K36me3, while regions of low antisense transcription are enriched for H3K79me2. Levels of all these histone modifications are affected by *PAF1* deletion. Therefore, deletion of *PAF1* and loss of Paf1C-dependent histone modifications may generate a chromatin landscape that promotes antisense transcription at some loci and represses it at others. Interestingly, in the *de novo* analysis, we observed enrichment of many transcripts oriented antisense to mRNA loci in the *paf1Δ trf4Δ* strain, relative to the *trf4Δ* strain, and found that Paf1 both positively and negatively regulates antisense transcript levels in *S. cerevisiae* (Figure 2B; Table S5).

Deeper analysis revealed that many of the antisense transcripts detected by the *de novo* analysis overlapped with previously annotated noncoding transcripts (Figure 2C), consistent with earlier studies showing that many noncoding transcripts are oriented antisense to genes (Xu *et al.* 2009; van Dijk *et al.* 2011; Schulz *et al.* 2013; Venkatesh *et al.* 2016). This suggests that a large portion of the ncRNA differential expression profile observed in the *paf1Δ trf4Δ* strain results from antisense transcription. To investigate the antisense transcriptional landscape further, we plotted sense and antisense transcript levels relative to the transcription start site (TSS) and transcription end site (TES) of protein-coding genes at which we detected an absolute change of 1.5-fold or greater in antisense transcription overlapping the gene (Figure S3A and Figure S3B). For both the antisense-downregulated class and the antisense-upregulated class, k-means clustering analysis revealed five clusters differing in the patterns and levels of antisense transcription relative to sense transcription. (Note that when these clusters are broken down by overlap with various ncRNA classes, no one cluster is dominated by an individual ncRNA class (Figure S3C, S3D, and Table S6)). A small number of protein-coding genes show an apparent anti-correlation between sense and antisense transcription in the *paf1Δ trf4Δ* mutant (cluster 1 in Figure S3A and cluster 5 in Figure S3B). However, for most genes experiencing a 1.5-fold or greater increase in antisense transcription, a clear relationship between antisense and sense transcript levels was not detected. This result agrees with previous work on antisense transcription (Murray *et al.* 2015). Further, a plot of sense and antisense transcript levels for all protein-coding genes suggests that antisense transcription is not governing the changes we detect in sense transcription for most genes (Figure S3E).

### Paf1 regulates transcript abundance at the transcriptional level

Paf1C has been shown to regulate both transcriptional and post-transcriptional processes at protein-coding genes (Porter *et al.* 2005; Van Oss *et al.* 2016; Fischl *et al.* 2017). To determine if changes in RNA levels detected by our tiling array analysis occurred at the transcriptional level, we compared our results to published NET-seq data (Harlen and Churchman 2017). Tiling array data comparing *paf1Δ* and *PAF1* strains or *paf1Δ trf4Δ* and *trf4Δ* strains were used to generate heatmaps for comparison to *paf1Δ* NET-seq data (Figure 3A). Overall, we observed similarity between the *paf1Δ trf4Δ* tiling array data and the *paf1Δ* NET-seq data, indicating that Paf1 is regulating the abundance of many transcripts, including unstable noncoding RNAs and mRNAs, through an effect on transcription (Figure 3A and 3B). However, our analysis also indicates, that at some genes, Paf1 is likely playing a post-transcriptional role. For example, for the majority of Paf1-stimulated protein-coding genes (73%), a decrease in steady state RNA levels in *paf1Δ* cells was reflected in reduced nascent transcript levels (Figure 3B). In contrast, a smaller fraction of Paf1-repressed genes (52%) showed a corresponding increase in NET-seq signal in the *paf1Δ* background (Figure 3B). Therefore, both positive and negative effects of Paf1 occur at the transcriptional level at many loci, but for protein-coding genes repressed by Paf1, a larger fraction appear to be post-transcriptionally regulated.

**Figure 3.**
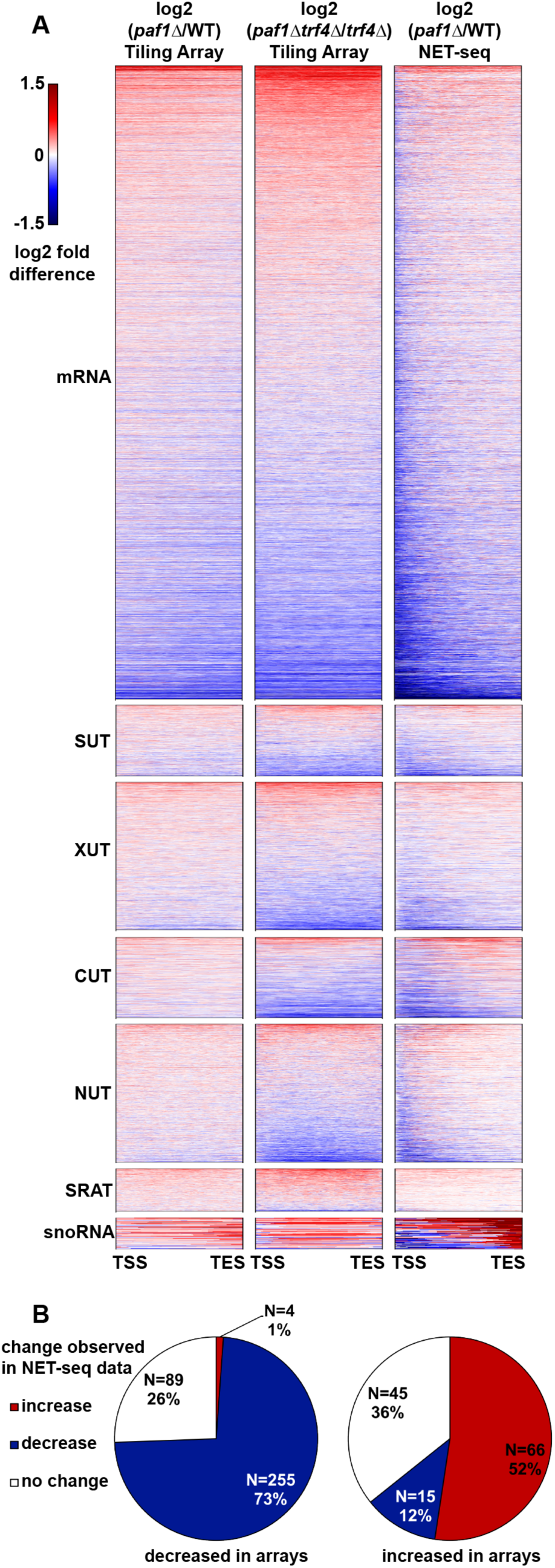
Paf1 regulates many of its target loci at the transcriptional level. (A) Heatmaps plotting log2 fold-change in transcript levels detected by tiling array for *paf1Δ* vs WT (KY1702 vs KY2276) and *paf1Δ trf4Δ* vs *trf4Δ* (KY2016 vs KY2012) as well as a *paf1Δ* vs WT NET-seq comparative analysis (Harlen and Churchman 2017). Previously annotated coding and non-coding transcripts were scaled so that each row in the heatmap represents a single transcript from transcription start site (TSS) to transcription end site (TES). (B) Pie charts showing the direction of change in NET-seq data (Harlen and Churchman, 2017) for mRNAs that increased or decreased expression by at least 1.5-fold in the *paf1Δ* vs WT comparison as measured by tiling array. Direction of change in NET-seq was determined by summing the reads in the first 500bp of protein-coding genes in both WT and *paf1Δ* NET-seq datasets and calculating a fold-change (1.5-fold cutoff).

One possible difference between Paf1-stimulated and Paf1-repressed genes is related to their level of expression in wild type cells. To investigate this possibility, we used ChIP-exo data from Van Oss et al. (2016) to analyze Pol II occupancy (Rpb3 subunit) at protein-coding genes with absolute expression changes of 1.5-fold or greater in a *paf1Δ* background as measured by our tiling array analysis. The Rpb3 ChIP-exo data indicate that, in general, Paf1-stimulated genes are more highly transcribed than Paf1-repressed genes (Figure S4). Similarly, Paf1 occupancy is higher at Paf1-stimulated genes compared to Paf1-repressed genes, consistent with the known association of Paf1C with Pol II. Since defects in Paf1C cause a disruption in telomeric silencing (Krogan *et al.* 2003; Ng *et al.* 2003a), we also analyzed the chromosomal locations of Paf1-regulated genes. Our analysis revealed broad chromosomal distribution of Paf1-stimulated and Paf1-repressed genes, in both *TRF4* and *trf4Δ* backgrounds, with no obvious bias toward telomeres (Figure S5).

### Paf1 stimulates the expression of phosphate homeostasis genes through a mechanism independent of its effects on individual histone modifications

Gene ontology analysis (Ashburner *et al.* 2000) revealed an enrichment in phosphate homeostasis genes among the genes where expression decreased upon deletion of *PAF1* in both the *TRF4* and *trf4Δ* backgrounds (Table 2). Given the wealth of information on the importance of chromatin structure in regulating genes in the phosphate regulon (Korber and Barbaric 2014), we explored the mechanism of Paf1 involvement at these genes. Our tiling array data show that many but not all genes activated by the Pho4 transcription factor (Zhou and O’Shea 2011) are downregulated in the absence of Paf1 (Figure 4A), arguing that the effects of Paf1 are unlikely to be due to a loss of Pho4 function. Consistent with this, *PHO4* transcript levels are not strongly affected in the *paf1Δ* strain (Figure 4A).

**Figure 4.**
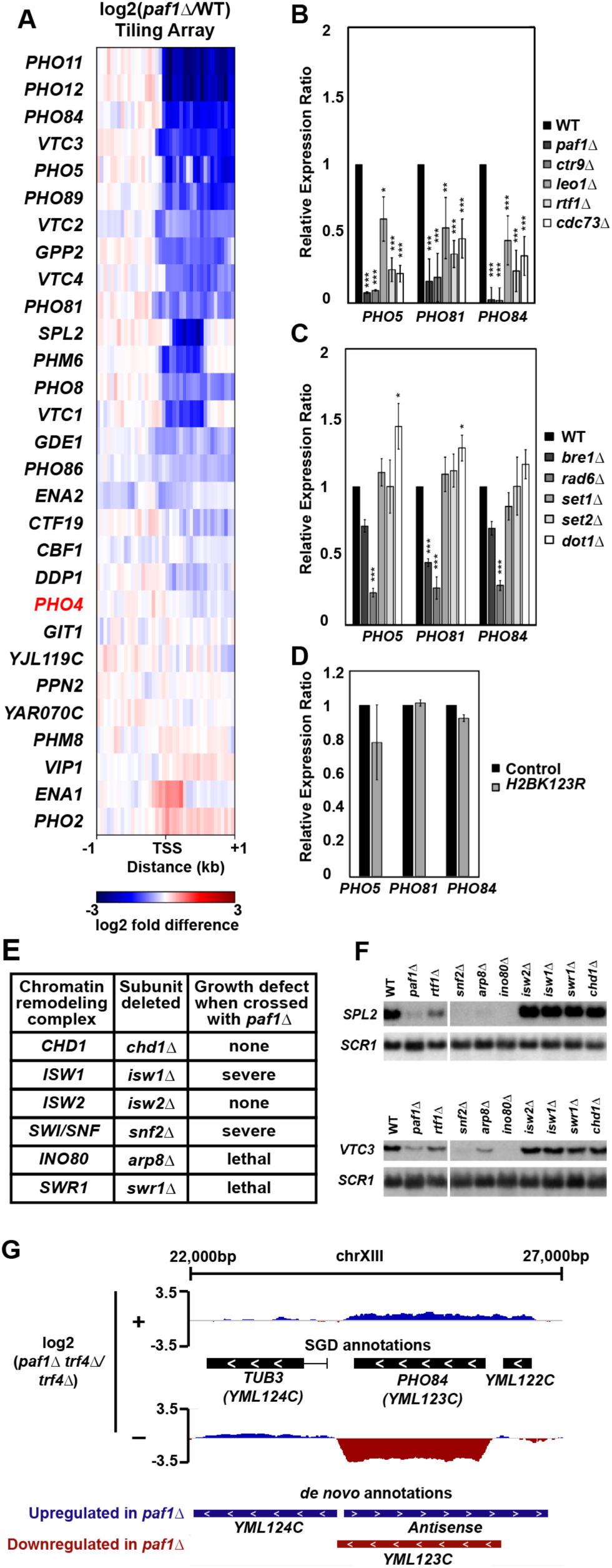
Paf1 positively regulates many phosphate homeostasis genes. (A) Heatmap of expression differences observed in a *paf1Δ* strain (KY1702) relative to a WT strain (KY2276) at Pho4-responsive genes (Zhou and O’Shea, 2011). (B-D) RT-qPCR analysis of phosphate gene expression in strains lacking (B) individual Paf1C subunits (KY1021, KY2271, KY2239, KY2243, KY2241 and KY2244), (C) histone modification enzymes (KY1683, KY2045, KY1952, KY938, KY914, KY934) or (D) H2B K123. In panel D, RNA levels in the H2B K123R mutant (KY2167) were compared to the appropriate WT control strain (KY2027). Relative expression ratio is calculated using primer efficiency, normalization to the RNA polymerase III transcript *SCR1* and a comparison to a WT strain (Pfaffl 2001). Error bars represent standard error of the mean and all statistically significant results are reported as asterisks (0.01 < p < 0.05 = *, 0.001 < p < 0.01 = **, 0 < p < 0.001 = ***). All *p* values were derived from a student’s t-test between the mutant strain and WT. (E) Cumulative data from crosses between a *paf1Δ* strain and strains deleted for chromatin remodeling factors. Following tetrad analysis of the following crosses, growth defects of double mutants were determined: *chd1Δ paf1Δ* = KY583 X KY804; *isw1Δ paf1Δ* = KY3464 X KY901; *isw2Δ paf1Δ* = KY884 X KY804; *snf2Δ paf1Δ* = KY508 X KY804; *arp8Δ paf1Δ* = KY3460 X KY804; *swr1Δ paf1Δ* = KY3462 X KY972. (F) Northern blot analysis of *SPL2* and *VTC3* RNA levels. Strains used in this analysis were KY292, KY802, KY508, KY3465, KY3461, KY884, KY3463, KY972 and KY632. *SCR1* serves as a loading control. (G) Genome browser view showing antisense transcription at the *PHO84* locus. The browser view shows smoothed differential expression tracks (log2(*paf1Δ trf4Δ / trf4Δ*), 160bp sliding window) with both SGD and *de novo* transcript annotations. Plus (+) and minus (−) symbols refer to DNA strand. The *PHO84* gene is oriented right to left.

To assess the contribution of individual Paf1C members to the expression of phosphate homeostasis genes, we performed RT-qPCR analysis of RNA isolated from *paf1Δ, ctr9Δ, cdc73Δ, rtf1Δ*, and *leo1Δ* strains. The RT-qPCR results generally agreed with our tiling array results. RNA levels for *PHO5, PHO81*, and *PHO84* were significantly decreased in the absence of any single Paf1C subunit with deletions of *PAF1* and *CTR9* causing the greatest effects (Figure 4B).

Given the prominent role of Paf1C in promoting transcription-coupled histone modifications, we asked if loss of these modifications could explain the gene expression changes we observed in the Paf1C mutant strains. To this end, we performed RT-qPCR assays on RNA prepared from strains lacking the H2Bub enzymes Rad6 and Bre1, the H3 K4 methyltransferase Set1, the H3 K36 methyltransferase Set2, or the H3 K79 methyltransferase Dot1 (Figure 4C). RNA levels for *PHO5, PHO81* and *PHO84* decreased in the *rad6Δ* and *bre1Δ* strains, which, like an *rtf1Δ* strain (Van Oss *et al.* 2016), are severely deficient in H2Bub. However, deletion of *PAF1* and *CTR9* had a substantially greater impact on the transcription of these genes than deletion of either *BRE1*, which encodes the ubiquitin ligase for H2B K123, or *RTF1*, which encodes the primary Paf1C determinant of H2Bub (Figures 4B and C). The larger effect of *rad6Δ* compared to *bre1Δ* suggests that, as a ubiquitin conjugating enzyme, Rad6 may play roles in *PHO* gene regulation beyond catalyzing H2Bub. In agreement with this, we observed only a slight decrease in *PHO5, PHO81*, and *PHO84* RNA levels in a H2B K123R mutant compared to the control strain (Figure 4D). Other than a slight, but statistically significant, upregulation of *PHO5* and *PHO81* in the *dot1Δ* strain, loss of individual H3 methyltransferases did not alter transcription of the *PHO* genes. Therefore, the loss of individual Paf1C-mediated histone modifications does not explain the strong reduction in phosphate homeostasis gene expression observed in *paf1Δ* and *ctr9Δ* mutants.

The absence of a clear effect of Paf1C-dependent histone modifications on *PHO* gene regulation prompted us to investigate other connections between Paf1 and chromatin. Previous work by Batta *et al.* (2012) showed reduced nucleosome occupancy within coding regions in a *paf1Δ* strain, and the importance of nucleosome occupancy changes for *PHO* gene expression have been well documented (Barbaric *et al.* 2007; Korber and Barbaric 2014). Therefore, we investigated genetic interactions between Paf1 and chromatin remodeling factors. Genetic crosses were performed between strains lacking Paf1 and strains mutated for the following chromatin remodeling factors: Chd1, Isw1, Isw2, Swi/Snf, Ino80, and Swr1 (Figure 4E). We observed synthetic lethality or severe synthetic growth defects in double mutants containing *paf1Δ* and a deletion of *SWR1, ISW1, SNF2* or *ARP8*, which encodes a subunit of the Ino80 complex. While the molecular basis for these genetic interactions is unclear, it is likely that Paf1C and chromatin remodeling factors regulate the expression of a shared group of genes. To test this idea for genes in the Pho4 regulon, we focused on two genes, *SPL2* and *VTC3*, known to be stimulated by Ino80 (Ohdate *et al.* 2003). Northern analysis revealed greatly reduced *VTC3* and *SPL2* expression in cells lacking *PAF1, SNF2, ARP8* or *INO80* (Figure 4F). Although deletion of *SWR1* or *ISW1* did not affect *SPL2* or *VTC3* mRNA levels, it is possible that other Paf1-dependent genes are regulated by these remodeling factors. Taken together, these data suggest that Paf1C and chromatin remodeling factors work in parallel to maintain gene expression levels required for cell viability and phosphate homeostasis.

A well-studied example of locus-specific antisense control of transcription occurs at the *PHO84* gene in yeast (Castelnuovo *et al.* 2013). At this locus, accumulation of an antisense transcript in an *rrp6Δ* strain leads to repression of the sense transcript through a mechanism dependent on particular histone modifications (Castelnuovo *et al.* 2014). In light of the changes in antisense RNAs detected in the *paf1Δ* background (Figure 2B and Figure S3), we examined our *de novo* differential expression data for evidence of Paf1-regulated antisense transcription at *PHO84* (Figure 4G). When comparing the *paf1Δ trf4Δ* mutant to the *trf4Δ* control strain, we observed increased antisense and decreased sense transcript levels across the *PHO84* gene. Interestingly, *PHO84* fell into one of the two clusters of genes for which an anti-correlation between sense and antisense transcription was observed in the *paf1Δ trf4Δ* mutant (Figure S3B, cluster 5). These data suggest that, at the *PHO84* gene, deletion of *PAF1* elevates antisense transcription and coordinately decreases sense transcription.

### Paf1 represses iron homeostasis gene expression in part through its role as a facilitator of H3 K36me3

Gene ontology analysis of genes that increased expression in *paf1Δ* strains revealed enrichment for genes in various iron-related processes (Table 2). As with the phosphate genes, an additional motivation for choosing iron homeostasis genes for follow-up experiments was the extent to which they have been characterized in the literature (Yamaguchi-Iwai *et al.* 2002; Rutherford and Bird 2004; Courel *et al.* 2005; Kaplan and Kaplan 2009; Cyert and Philpott 2013). Our tiling array analysis of genes that are normally activated by the Aft1 and Aft2 transcription factors in iron-limiting conditions (Cyert and Philpott 2013) revealed that many but not all of these genes are repressed by Paf1 in iron-replete media (Figure 5A).

**Figure 5.**
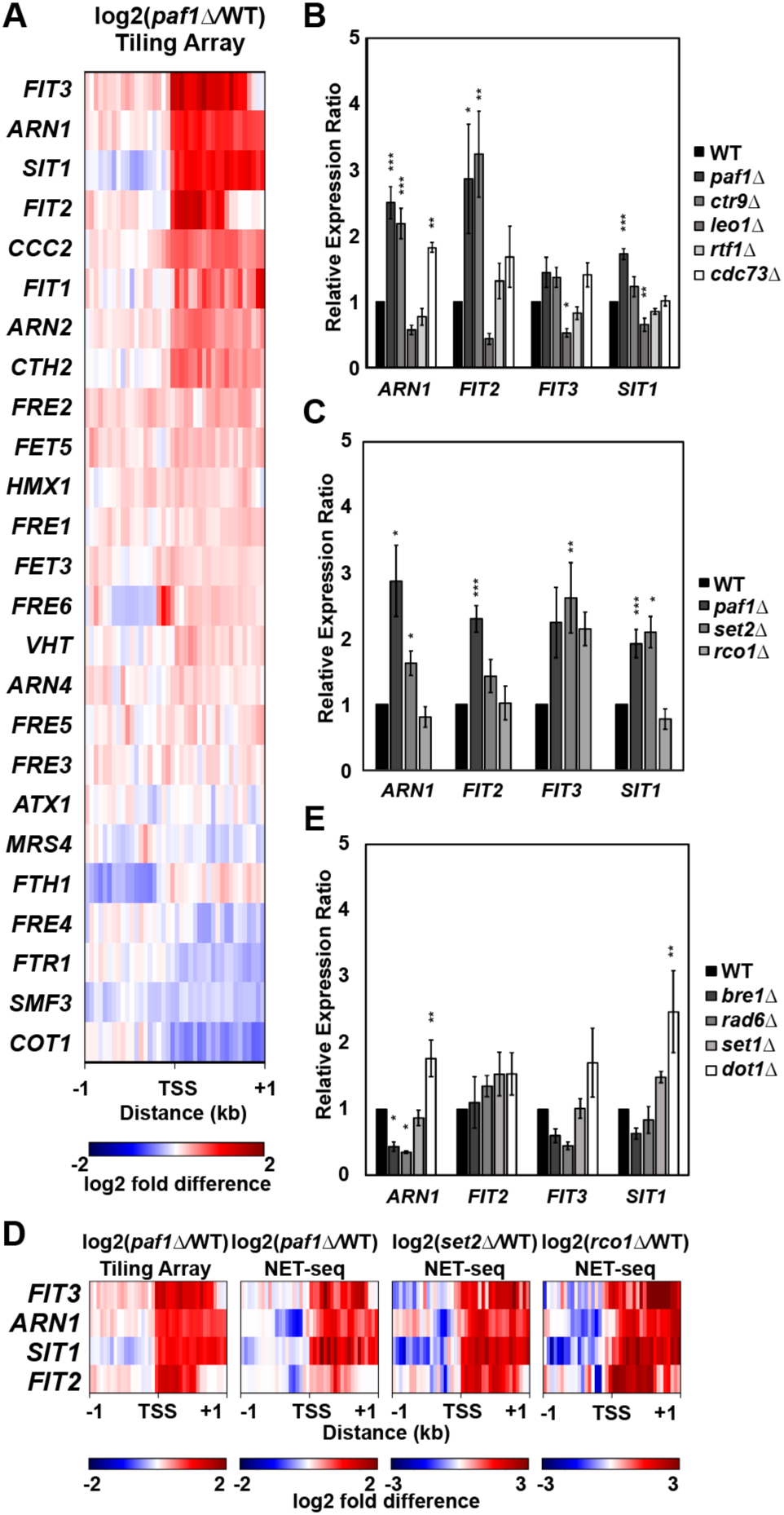
Paf1 represses iron homeostasis genes. (A) Heatmap of expression differences observed in a *paf1Δ* strain (KY1702) relative to a WT strain (KY2276) at Aft1 and Aft2 responsive genes involved in maintenance of iron homeostasis (Cyert and Philpott 2013). (B-C) RT-qPCR analysis of the indicated genes in strains lacking (B) individual Paf1C subunits (KY1021, KY2271, KY2239, KY2243, KY2241 and KY2244) or (C) genes in the Set2/Rpd3S pathway (KY307, KY914, KY1702 and KY1235). Calculation of the relative expression ratio and statistical testing were performed as in Figure 4. (D) Heatmaps of expression differences between mutant yeast strains and their respective WT strains in tiling array (this study) and NET-seq (Churchman and Weissman 2011; Harlen and Churchman 2017) datasets. (E) RT-qPCR results for iron homeostasis genes in strains lacking enzymes that catalyze Paf1C-associated histone modifications (KY1683, KY2045, KY1952, KY938, and KY934).

To further investigate this subset of genes, we performed RT-qPCR analysis on RNA isolated from strains lacking individual Paf1C members (Figure 5B). A reproducible increase in expression was observed for *ARN1, FIT2, FIT3* and *SIT1* in *paf1Δ* and *ctr9Δ* strains. With the exception of *SIT1*, deletion of *CDC73* also led to derepression of these genes. In contrast, whereas Paf1, Ctr9, and Cdc73 repress the transcription of *ARN1, FIT2*, and *FIT3*, Leo1 appears to play a stimulatory role at these genes, while Rtf1 has little effect. Together, these data demonstrate that individual Paf1C subunits differentially regulate iron-responsive genes.

The Set2 histone methyltransferase catalyzes H3 K36me3, a modification that is dependent on Paf1 and Ctr9 (Chu *et al.* 2007). This epigenetic mark leads to the activation of the Rpd3S histone deacetylase complex and inhibition of histone exchange, generating a repressed chromatin state (Carrozza *et al.* 2005; Joshi and Struhl 2005; Keogh *et al.* 2005; Govind *et al.* 2010; Churchman and Weissman 2011; Venkatesh *et al.* 2012, 2016; Kim *et al.* 2016). In the *set2Δ* strain, RNA levels for *FIT3* and *SIT1* increased to those observed in the *paf1Δ* strain, suggesting that Paf1 represses these genes through stimulating H3 K36me3 (Figure 5C). Further, NET-seq data (Churchman and Weissman 2011; Harlen and Churchman 2017) indicate that the increase in steady state mRNA levels for *FIT3* and *SIT1* in *paf1Δ* and *set2Δ* strains is associated with an increase in transcription (Figure 5D).

For two other genes, *ARN1* and *FIT2*, the level of derepression observed in the *paf1Δ* strain was significantly higher than that observed in the *set2Δ* strain, despite evidence from NET-seq data (Figure 5D; Churchman and Weissman 2011; Harlen and Churchman 2017) that loss of Set2 strongly increases transcription of these genes. Similarly, with the exception of *FIT3*, steady state mRNA levels in a strain lacking the Rpd3S subunit Rco1 did not reflect the increase in transcription detected by NET-seq (Figure 5C and 5D). One likely explanation for the difference between the steady state mRNA measurements (Figure 5C) and the nascent transcript data (Figure 5D) is that mRNA levels for the iron-responsive genes are post-transcriptionally regulated, possibly through a degradation pathway that involves Paf1. This conclusion is in line with observations made through our *de novo* analysis (Figure 3B), which indicated a post-transcriptional role for Paf1 at genes where it negatively regulates mRNA levels, and with previous descriptions of RNA degradation pathways that target mRNAs in the iron regulon (Lee *et al.* 2005; Puig *et al.* 2005).

In addition to the Set2/Rpd3S pathway, we tested other histone modifiers for a role in iron gene repression by examining *ARN1, FIT2, FIT3* and *SIT1* expression in *bre1Δ, rad6Δ, set1Δ*, and *dot1Δ* strains by RT-qPCR (Figure 5E). With the exception of the *dot1Δ* mutation, which elevated *ARN1* and *SIT1* transcript levels, none of these mutations led to a significant derepression of the iron genes. Taken together, these results suggest that Paf1 represses expression of genes in the iron regulon by inhibiting transcription, most likely by facilitating histone marks such as H3 K36me3, and by influencing RNA stability.

### *FET4* is differentially regulated by Paf1 and upstream CUT transcription

To investigate the interplay between Paf1 and noncoding DNA transcription on a protein-coding gene in the iron regulon, we focused on the *FET4* gene, which encodes a low affinity iron transporter in *S. cerevisiae* (Dix *et al.* 1994). Two CUTs have been annotated upstream of the *FET4* coding region (Xu *et al.* 2009; Raupach *et al.* 2016) (Figure 6A). We hypothesized that *CUT 794/793* transcription regulates *FET4* transcription possibly in a *PAF1-*dependent manner. To test this, we generated strains containing a transcription termination sequence (TTS) upstream of the *FET4* gene positioned to stop transcription of both upstream CUTs in wild type, *paf1Δ, trf4Δ* and *paf1Δ trf4Δ* backgrounds.

**Figure 6.**
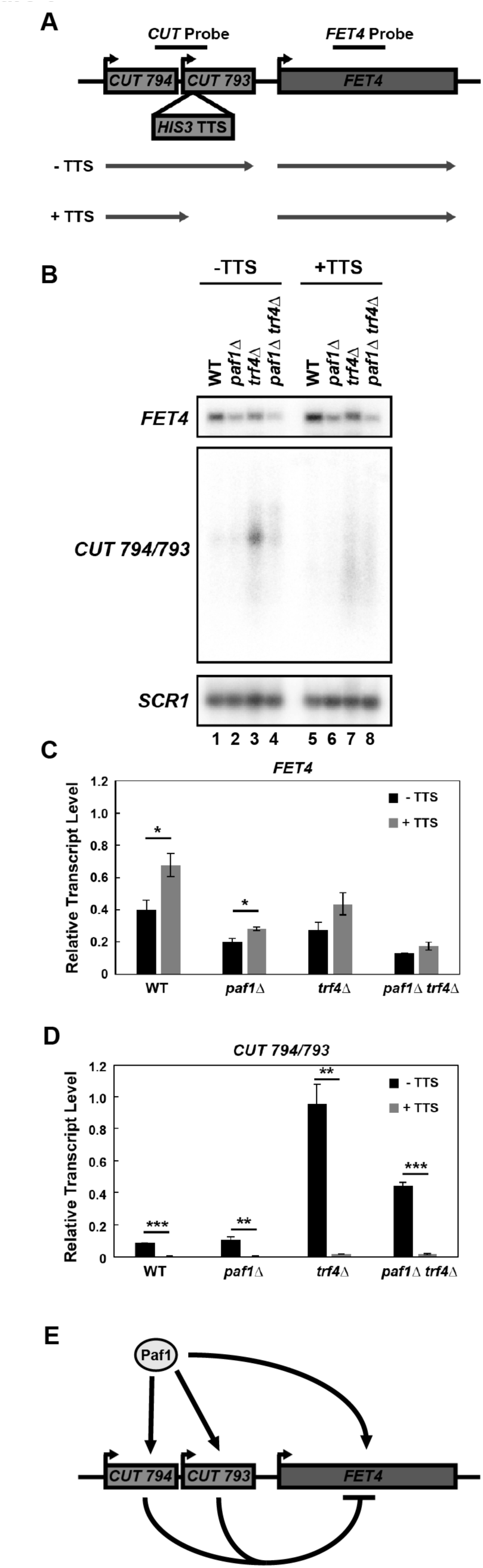
The *FET4* locus is regulated by Paf1 and transcription of ncDNA upstream of the coding region. (A) Diagram of the *FET4* locus and the position of a transcription termination sequence (*HIS3* TTS) inserted 400 bp upstream of the *FET4* start codon to block *CUT 794/793* transcription. (B) Northern analysis of *FET4* mRNA, *CUT 794/793* and *SCR1* RNA (loading control) from WT, *paf1Δ, trf4Δ*, and *paf1Δ trf4Δ* strains without the inserted TTS (KY2276, KY1702, KY2012, KY2016) or with the TTS (KY3466, KY2846, KY2851, KY2845). (C-D) Quantification of northern blot results for *FET4* and *CUT 794/793* normalized to *SCR1*. Error bars represent standard error of the mean and all statistically significant results are reported as asterisks that represent *p* values from students t-test as in Figure 4. (E) Diagram of the observed effects of *PAF1* and *CUT794/793* at the *FET4* locus.

Northern analysis showed that deletion of *PAF1* reduced *FET4* transcript levels (Figure 6B top blot, compare lanes 1 and 2 and lanes 3 and 4; Figure 6C) and *CUT794/793* levels (Figure 6B middle blot, compare lanes 3 and 4; Figure 6D). When the TTS was inserted upstream of *FET4* (+TTS), CUT levels decreased and *FET4* transcript levels increased in all conditions tested, suggesting that transcription of the upstream CUT inhibits expression of the coding region (Figures 6B-D). This is reminiscent of the inhibitory effect of noncoding transcription upstream of the well-studied *SER3* gene (Martens *et al.* 2004). Interestingly, even when CUT transcription was blocked, *FET4* transcript levels were reduced in the *paf1Δ* background. These data suggest that CUT transcription upstream of the *FET4* promoter negatively regulates transcription of the *FET4* gene and that Paf1 has a dual role in regulating *FET4* by stimulating expression of both the ORF and the inhibitory *CUTs 794/793* (Figure 6E).

## DISCUSSION

The many roles of Paf1C and the pleiotropic phenotypes conferred by deletion of individual Paf1C subunits (Betz *et al.* 2002) suggest that this conserved transcription elongation complex regulates the expression of many genetic loci. While previous studies focused on the regulation of mRNAs (Shi *et al.* 1996; Penheiter *et al.* 2005; Porter *et al.* 2005; Batta *et al.* 2011; van Bakel *et al.* 2013; Cao *et al.* 2015; Chen *et al.* 2015; Yang *et al.* 2016; Fischl *et al.* 2017; Harlen and Churchman 2017), here we sought to identify the full cohort of Paf1-regulated transcripts, both coding and noncoding, by exploiting a genetic background deficient in the TRAMP/exosome-dependent RNA degradation pathway and by performing *de novo* transcript identification analyses. Our high-resolution tiling array experiments revealed differential expression of transcripts in all Pol II transcribed RNA classes in strains deleted for *PAF1.* A comparison of our *paf1Δ trf4Δ* tiling array data with published *paf1Δ* NET-seq (Harlen and Churchman 2017) data demonstrated that Paf1 regulates many coding and noncoding RNAs at the transcriptional level and that the presence of a functional TRAMP complex obscures many of these transcriptional effects.

Our study revealed both positive and negative roles of Paf1 in regulating noncoding RNA levels. For many transcripts in the CUT, NUT, XUT, and SUT classes, Paf1 stimulates their expression. For two other important classes of noncoding RNAs, snoRNAs and SRATs, Paf1 functions primarily as a negative regulator. The elevation in SRAT expression was not unexpected given the importance of Paf1 for H3 K36me3 (Chu *et al.* 2007), a mark important for the maintenance of a repressive chromatin environment. In our previous work, we showed that Paf1 is important for snoRNA termination (Sheldon *et al.* 2005; Tomson *et al.* 2011, 2013). We recapitulate those results here and identify additional snoRNA loci that exhibit transcription termination defects in the absence of Paf1 (Figure 1H). These results, together with our finding that Paf1 impacts the transcription of many CUTs and NUTs, extends the functional connections between Paf1 and the machinery that terminates and processes these noncoding RNAs, including the NNS machinery and the nuclear exosome. Similar to a *paf1Δ* strain, snoRNA 3’ ends are extended in *rrp6* and *nrd1* mutants, which lack subunits of the nuclear exosome and NNS, respectively (Schulz *et al.* 2013; Fox *et al.* 2015). In contrast, while NUTs and CUTs are elevated in *nrd1* and *rrp6* mutants, levels of many of these unstable noncoding RNAs are decreased in strains deleted for *PAF1*. The reduced levels of these RNAs in *paf1Δ* strains are likely due, at least in part, to the stimulatory effect Paf1 has on their transcription (Figure 3).

By performing a *de novo* differential expression analysis of our tiling array data, we uncovered effects of Paf1 on antisense transcription (Figure 2B). Interestingly, many of the histone modifications promoted by Paf1C are reduced in regions experiencing higher levels of antisense transcription, but still others are present at high levels in these same regions (Murray *et al.* 2015). The loss of Paf1C-promoted histone modifications may therefore contribute to changes in antisense transcription (Castelnuovo *et al.* 2014; Murray *et al.* 2015). Indeed, we found instances of both increased and decreased antisense transcription in our *paf1Δ* strains. Although some global anticorrelation exists between sense and divergent antisense transcription initiating from nucleosome depleted regions (Xu *et al.* 2009; Churchman and Weissman 2011), antisense transcription does not universally correlate or anticorrelate with sense transcription (Murray *et al.* 2015). Our results agree with this observation (Figure S3E), but also point to a small subset of genes where sense and antisense transcription appear to be anticorrelated when *PAF1* is deleted (Figure S3A cluster 1 and S3B cluster 5).

One gene that fits into this category is *PHO84* (Figure S3B cluster 5; Figure 4G). Our data suggest a role for Paf1 in preventing antisense transcription at *PHO84* independently of its functional connections with the TRAMP/exosome pathway, as we detect higher antisense and lower sense transcript levels in the *paf1Δ trf4Δ* strain relative to the *trf4Δ* strain (Figure 4G). The ability to detect changes in antisense transcription was enhanced by the absence of Trf4. This observation agrees with studies on *PHO84* and other genes, which showed elevated antisense transcription in the absence of Rrp6 (Castelnuovo *et al.* 2013). With respect to *PHO84*, the mechanism by which Paf1 facilitates sense and represses antisense transcription remains undefined. Although previous studies showed that *set1Δ* strongly upregulates *PHO84* sense transcription in the presence of *RRP6* (Castelnuovo *et al.* 2013) and that Paf1C is required for Set1-dependent H3 K4 methylation (Krogan *et al.* 2003; Ng *et al.* 2003b), our strand-specific tiling array data showed that *paf1Δ* strongly downregulates *PHO84* sense transcription (Figure 4A). Similarly, our results do not ascribe the stimulatory effect of Paf1 on *PHO5* and *PHO81* to any single Paf1C-dependent histone modification or an obvious change in antisense transcription; however, it remains possible that the individual modifications function redundantly in promoting the expression of these genes.

At many genes that are normally induced in iron-limiting conditions, Paf1 plays a repressive role under iron-replete conditions. For the four strongly upregulated genes examined, deletion of *PAF1* increased steady state RNA levels as well as nascent transcript levels, arguing that Paf1 is controlling the transcription of these genes. The increase in transcription in the *paf1Δ* strain correlates with a decrease in Set2 function, as shown by NET-seq data (Churchman and Weissman 2011; Harlen and Churchman 2017). Indeed, in our tiling array experiments, we detected a global increase in SRAT transcription in a *paf1Δ* strain (Figure 1D and 2A). Interestingly, when comparing steady state RNA levels and nascent transcript levels, we noted an apparent post-transcriptional effect of Paf1 (Figures 3B, 5C and 5D). With respect to the iron metabolism genes, we saw a strong overlap between Paf1-repressed mRNAs and Rnt1-repressed mRNAs (Lee *et al.* 2005). Rnt1 is a double-stranded RNA endonuclease that cleaves RNAs with a particular stem-loop structure (Chanfreau *et al.* 2000) and, in iron replete conditions, executes an RNA degradation pathway for mRNAs that encode iron uptake proteins (Lee *et al.* 2005). While other explanations are possible, the overlap between Paf1- and Rnt1-repressed mRNAs suggests that the post-transcriptional role of Paf1 at iron regulon genes may involve a functional interaction with Rnt1. A recent study showed that the rate of transcription elongation can influence the folding and processing of histone pre-mRNAs (Saldi *et al.* 2018), raising the possibility that deletion of *PAF1* might alter the rate of elongation in a way that affects the folding of substrates for Rnt1 or another RNA processing factor. Together, our results suggest that through stimulating Set2-mediated H3 K36 methylation, Paf1 represses genes in the iron regulon, but has an additional role in reducing the stability of these mRNAs.

Numerous examples of protein-coding gene regulation by ncDNA transcription or ncRNAs have been observed (Castelnuovo and Stutz 2015). Adding to this body of evidence we investigated the regulatory mechanisms governing expression of *FET4*, which encodes a low affinity iron transporter. Our analysis indicates that *FET4* is regulated by the expression of upstream CUTs and by Paf1. Insertion of a transcription termination sequence upstream of *FET4* decreased *CUT794/793* levels and increased *FET4* transcription. Deletion of *PAF1* reduced both *CUT794/793* and *FET4* transcript levels. Together with these targeted experiments, our tiling array results on the *paf1Δ* strain also revealed a stimulatory effect of Paf1 on *FET4* mRNA levels. However, we note that our tiling array analysis of the *paf1Δ trf4Δ* strain indicated that, in some circumstances, Paf1 can repress *FET4* expression. Previous studies have shown that genetic background and growth conditions can influence the levels of *FET4* mRNA and the noncoding RNAs adjacent to or overlapping *FET4* (*CUT794/793* and the *SUT322* antisense ncRNA) (Xu *et al.* 2009). Since our tiling array and northern blotting experiments used RNA from yeast grown on separate days, it is possible that slight differences in media may be responsible for differences in expression dynamics at the *FET4* locus, highlighting the intricacies of the regulatory system operating at this gene. Collectively, our results add to the interesting list of telomere-proximal metal-responsive genes under the control of noncoding transcription (Toesca *et al.* 2011).

The complexity of the transcription process and its regulation by chromatin provides numerous opportunities for multifunctional transcription factors, like Paf1C, to regulate gene expression. Our study reveals genome-wide effects of Paf1 on both coding and noncoding RNAs and provides mechanistic explanations for its diverse effects on specific classes of protein-coding genes. An understanding of the locus-specific effects of Paf1C will be an important step in elucidating the numerous connections of this complex to gene expression changes that cause human disease (Tomson and Arndt 2013; Karmakar *et al.* 2018).

## ACKNOWLEDGEMENTS

We thank Margaret Shirra, Elizabeth Hildreth, Christine Cucinotta, Matthew Hurton, and Paul Cantalupo for helpful discussions. We are grateful to Frank Pugh and Stirling Churchman, and members of their labs, for assistance with analysis of published genomic data, and to Chenchen Zhu and Lars Steinmetz for advice on building a probe annotation file for the tiling array analysis. This research was supported by the National Institutes of Health Grant R01 GM52593 to K.M.A., CRC funds to C.N., and funds from the University of Pittsburgh to M.L. M.A.E. is supported by a predoctoral fellowship from the NIH (F31GM129917).

## SUPPLEMENTAL FIGURE LEGENDS

**Figure S1.**
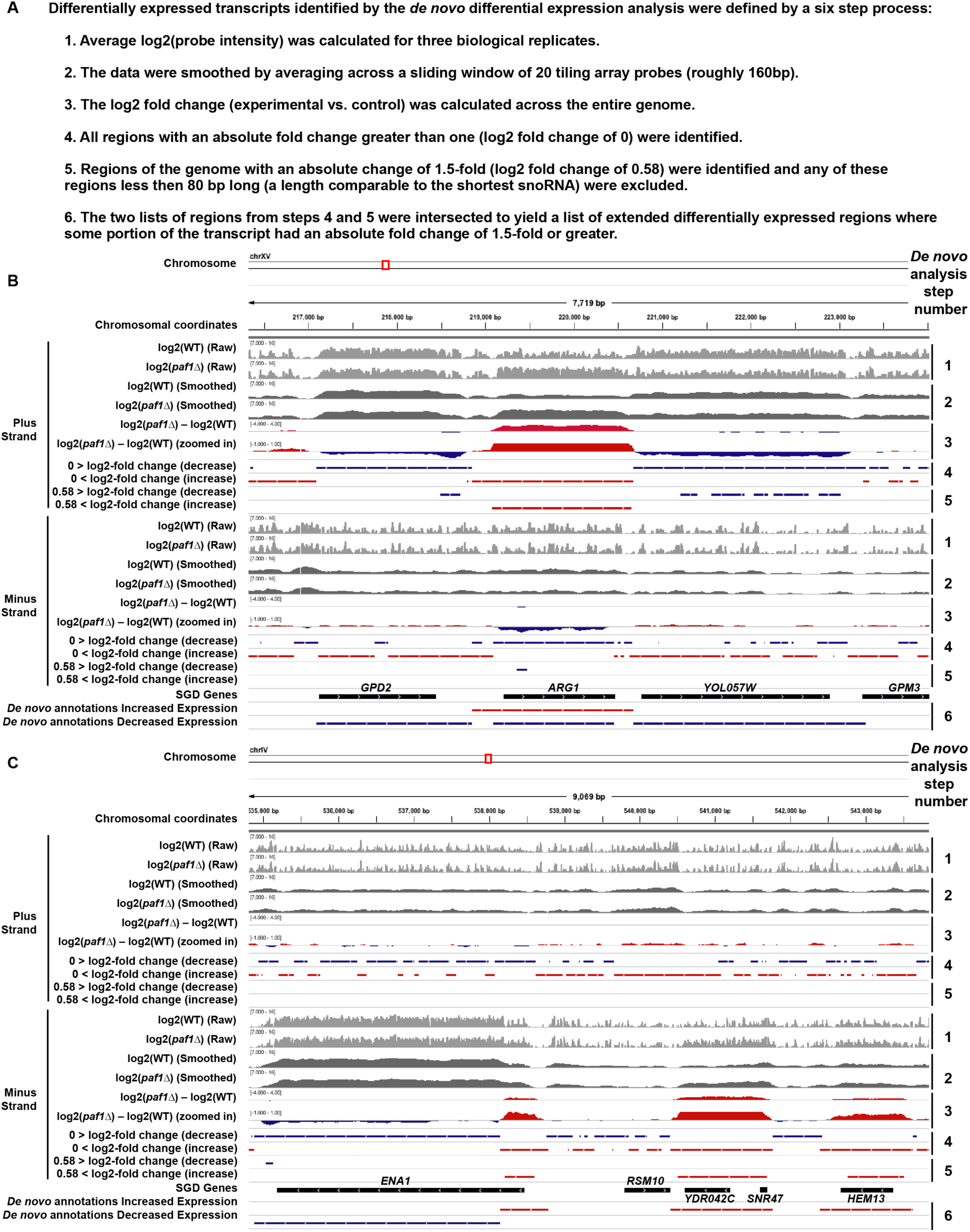
Examples of the steps taken in the *de novo* differential expression analysis. (A) List of steps taken to identify differentially expressed transcripts using the *de novo* analysis. (B) Genome browser tracks generated using IGV showing step by step how differentially expressed transcripts are identified by the *de novo* analysis. Browser tracks show tiling array data from the *paf1Δ* and WT datasets. Numbers corresponding to steps listed in A are shown on the right. (C) Same as A at a different genomic location.

**Figure S2.**
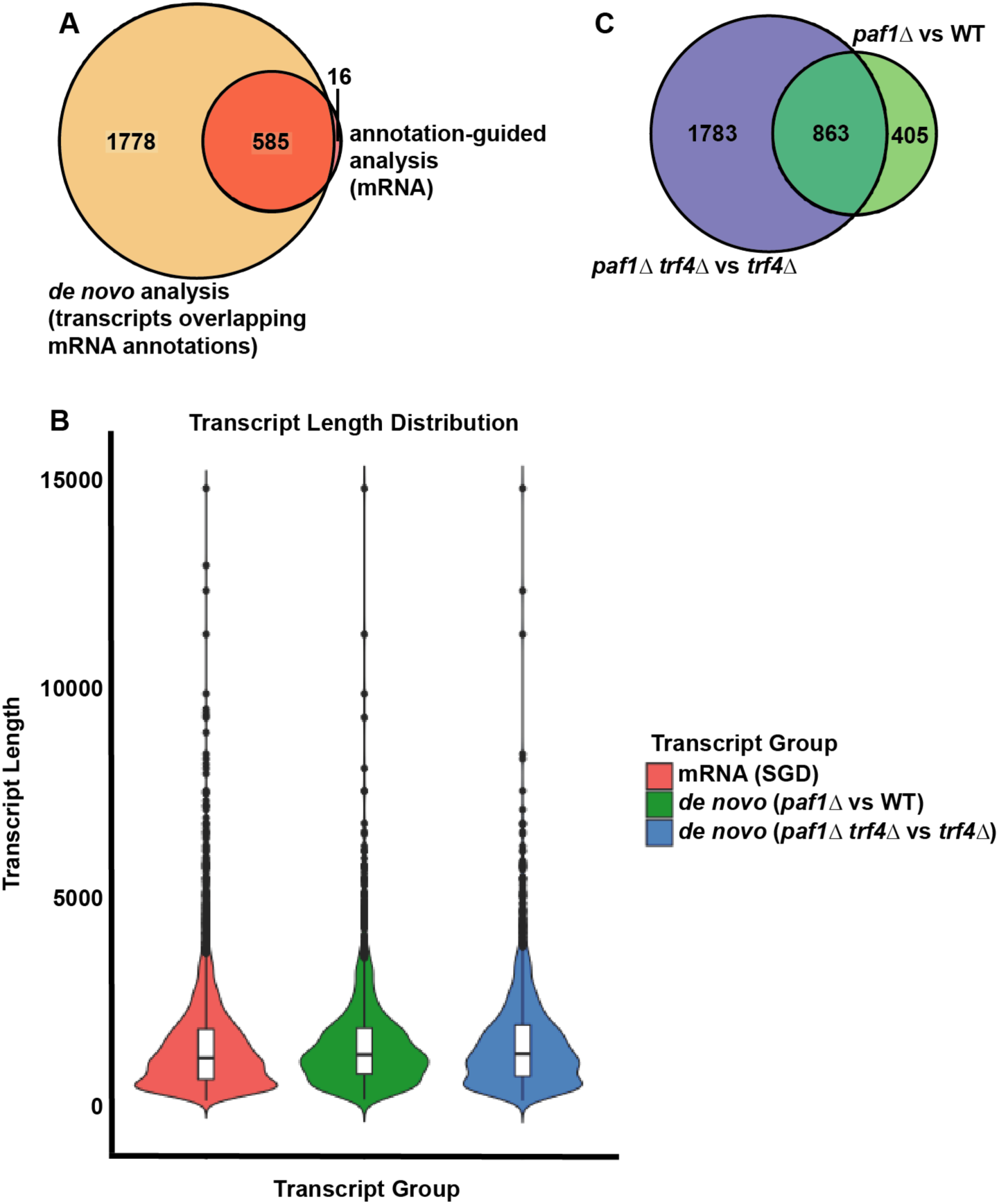
Comparison of the annotation-guided and *de novo* differential expression analyses. (A) Venn diagram comparing differentially expressed transcripts that overlap with mRNA annotations in the *paf1Δ trf4Δ* strain (KY2016) identified by *de novo* analysis (1.5-fold cutoff and length greater than 80bp) and differentially expressed mRNAs identified in our annotation-guided analysis (1.5-fold cutoff). When more than one differentially expressed transcript, as identified by *de novo* analysis, overlapped with the same mRNA, the overlap was only counted once in the intersecting region of the Venn diagram. (B) Violin plots showing the distribution of transcript lengths for mRNA annotations in SGD and the *de novo* annotations from this study. (C) Venn diagram showing overlap between all differentially expressed transcripts identified by *de novo* analysis of *paf1Δ* and *paf1Δ trf4Δ* strains (KY1702 and KY2016, respectively).

**Figure S3.**
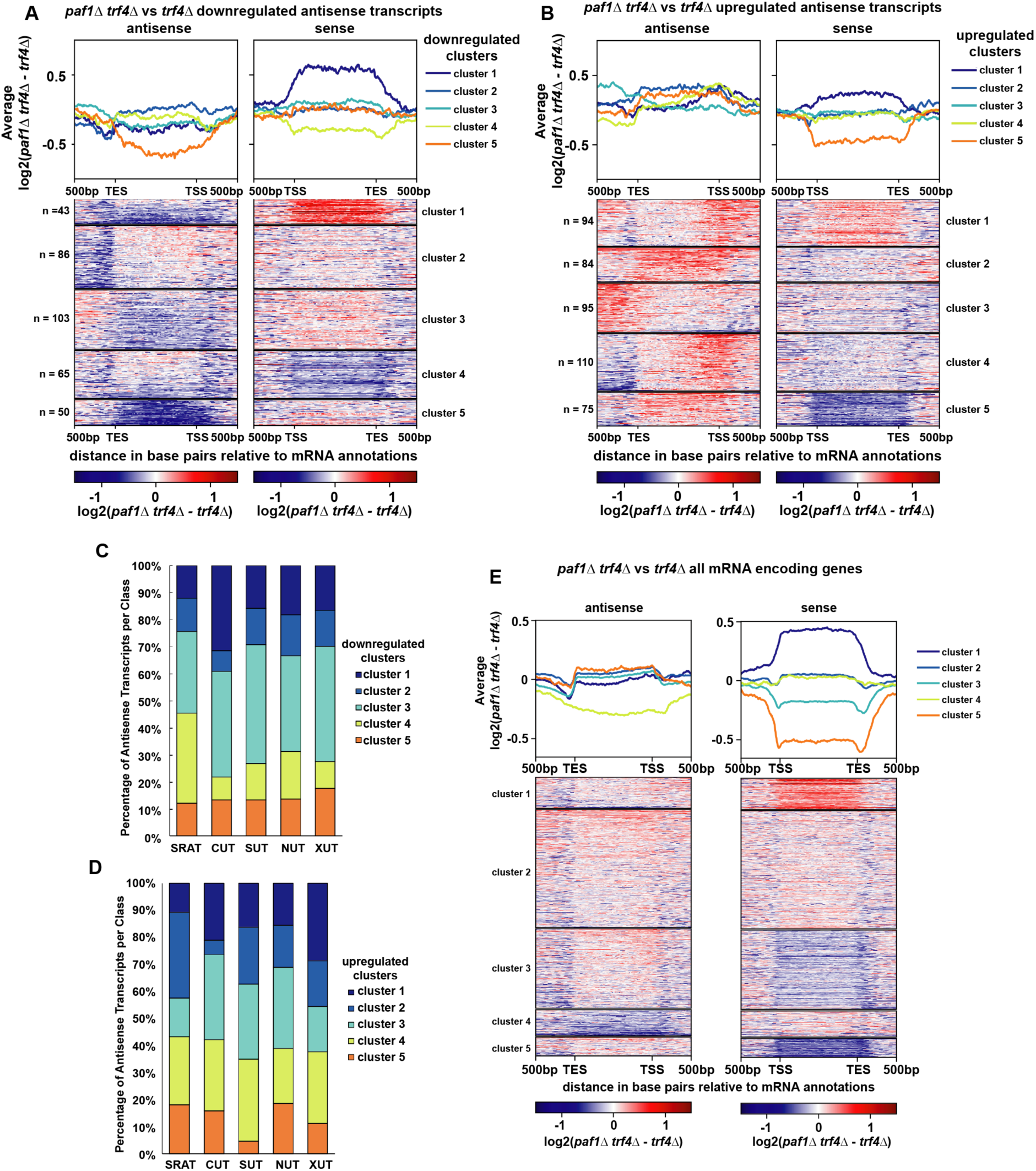
Analysis of antisense transcription in the *paf1Δ trf4Δ* mutant. (A and B) Heatmaps and average gene profiles for clusters generated by k-means clustering of the sense and antisense strands of protein-coding regions that experience changes in antisense transcription in the *paf1Δ trf4Δ* vs *trf4Δ* shown in Figure 2B. Protein-coding genes with decreased and increased antisense transcription are shown in panels A and B, respectively. (C and D) Vertically stacked bar graphs showing the percentage of regions antisense to mRNAs overlapping with various noncoding transcript classes. Clusters were taken from the analyses in A and B. (E) Heatmaps and average gene profiles of tiling array data (log2(*paf1Δ trf4Δ*) – log2(*trf4Δ*)) on the sense and antisense strand of all protein-coding genes. These data are scaled over the gene body and an additional 500 bp upstream and downstream are shown. These data were separated into clusters using k-means clustering.

**Figure S4.**
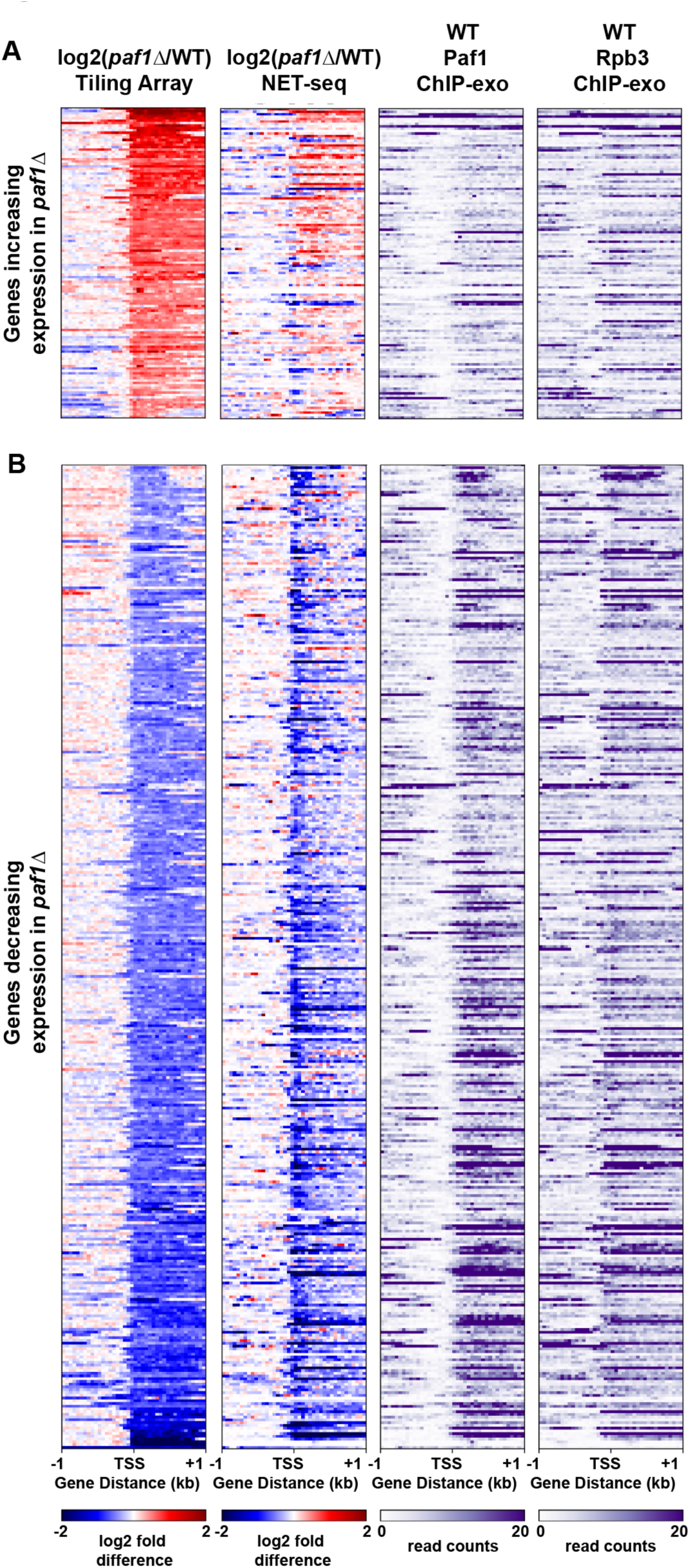
Differentially regulated protein-coding genes shown by tiling array, NET-seq, and ChIP-exo across three different studies. (A) Heatmaps of genes that increased expression by 1.5-fold or more in the *paf1Δ* strain relative to WT. (B) Heatmaps of genes that decreased expression by at least 1.5-fold in the *paf1Δ* strain relative to WT. Gene lists were determined by selecting genes that decreased or increased expression by at least 1.5-fold in the tiling array data presented here. Heatmaps were sorted by the tiling array data values. NET-seq data were taken from (Harlen and Churchman 2017) and ChIP-exo data were taken from (Van Oss *et al.* 2016). All heatmaps are plotted relative to the transcription start site (TSS). Regions 1kb upstream and downstream of the TSS are shown.

**Figure S5.**
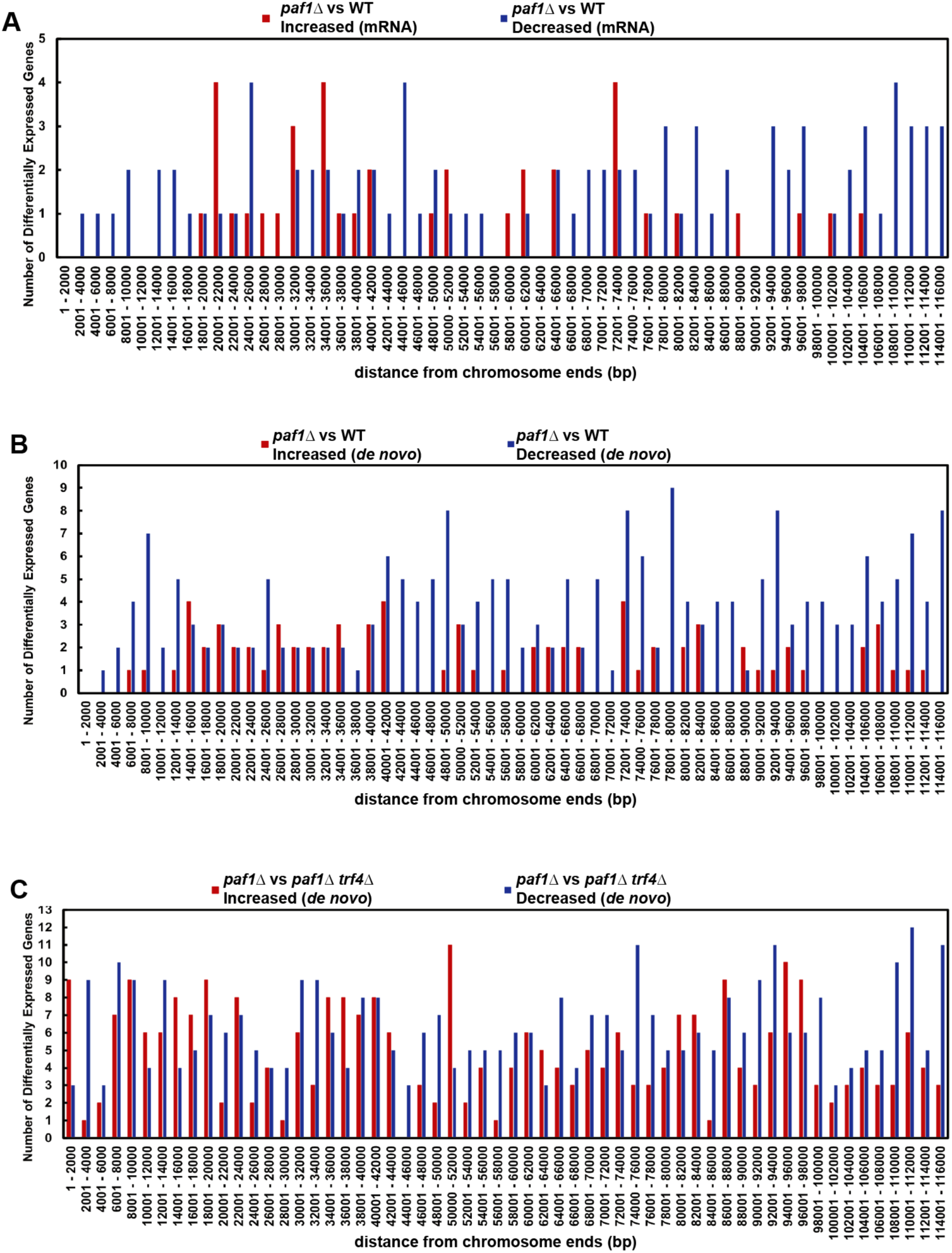
The positions of Paf1-regulated transcripts are not strongly biased toward the telomere. Non-overlapping bins of 2000 bp were generated working in toward the centromere from both ends of each yeast chromosome. Bins were intersected with transcript annotations to generate count tables. Bar graphs show the number of transcripts within each bin that increased (red) or decreased (blue) in the *paf1Δ* background. (A) Counts of differentially expressed mRNAs in a *paf1Δ* strain relative to WT. (B) Counts of differentially expressed transcripts identified in the *de novo* analysis comparing *paf1Δ* to WT. (C) Counts of differentially expressed transcripts identified in the *de novo* analysis comparing *paf1Δ trf4Δ* to *trf4Δ.*

**Table S1.**
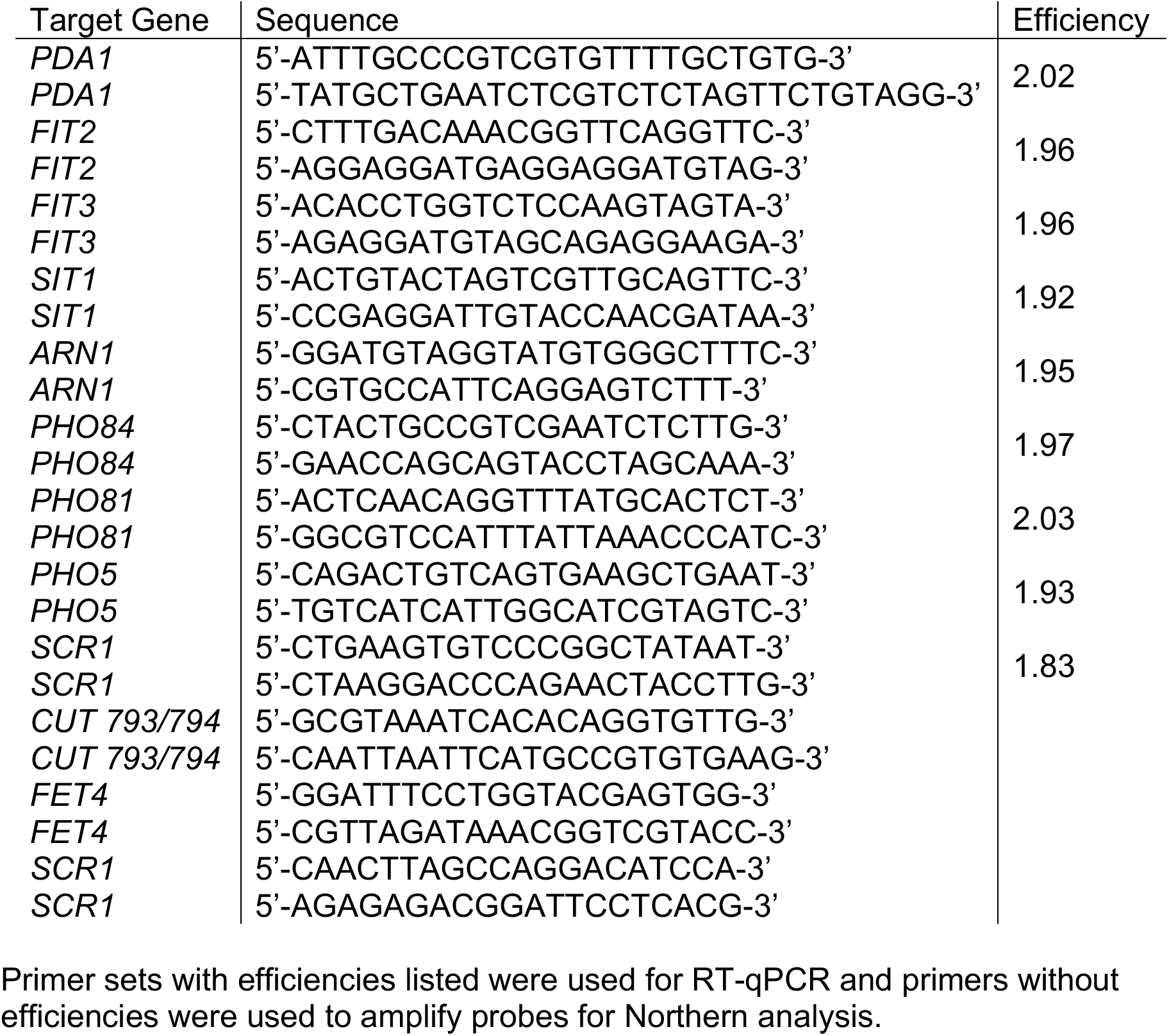
Primers for RT-qPCR and Northern probe generation

**Table S2.**
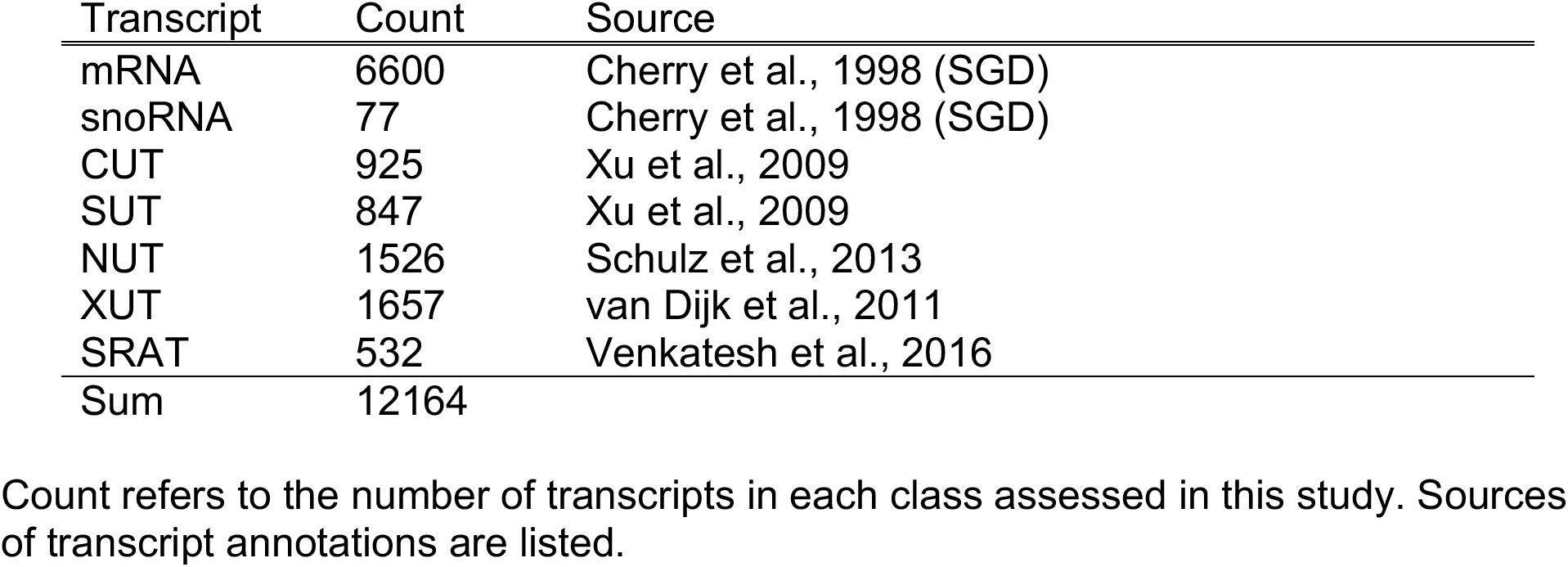
Transcript counts

| Transcript | Count | Source |
| --- | --- | --- |
| mRNA | 6600 | Cherry et al., 1998 (SGD) |
| snoRNA | 77 | Cherry et al., 1998 (SGD) |
| CUT | 925 | Xu et al., 2009 |
| SUT | 847 | Xu et al., 2009 |
| NUT | 1526 | Schulz et al., 2013 |
| XUT | 1657 | van Dijk et al., 2011 |
| SRAT | 532 | Venkatesh et al., 2016 |
| Sum | 12164 |  |

Count refers to the number of transcripts in each class assessed in this study. Sources of transcript annotations are listed.

**Table S3.** Summary of differential expression results obtained from the annotation-guided analysis of the tiling array data

| Transcript Class | *paf1Δ vs WT* |  |  |  |  |  |
| --- | --- | --- | --- | --- | --- | --- |
|  | Increase |  | Decrease |  | Sum |  |
|  | Count | Class Percent | Count | Class Percent | Count | Class Percent |
| mRNA | 126 | 1.9 | 348 | 5.3 | 474 | 7.2 |
| snoRNA | 12 | 15.6 | 1 | 1.3 | 13 | 16.9 |
| CUT | 8 | 0.9 | 2 | 0.2 | 10 | 1.1 |
| SUT | 4 | 0.5 | 3 | 0.4 | 7 | 0.9 |
| NUT | 9 | 0.6 | 5 | 0.3 | 14 | 0.9 |
| XUT | 10 | 0.6 | 5 | 0.3 | 15 | 0.9 |
| SRAT | 2 | 0.4 | 0 | 0.0 | 2 | 0.4 |
| Sum | 171 |  | 364 |  | 535 |  |

| Transcript Class | *paf1Δ trf4Δ vs trf4Δ* |  |  |  |  |  |
| --- | --- | --- | --- | --- | --- | --- |
|  | Increase |  | Decrease |  | Sum |  |
|  | Count | Class Percent | Count | Class Percent | Count | Class Percent |
| mRNA | 281 | 4.3 | 320 | 4.8 | 601 | 9.1 |
| snoRNA | 21 | 27.3 | 2 | 2.6 | 23 | 29.9 |
| CUT | 14 | 1.5 | 86 | 9.3 | 100 | 10.8 |
| SUT | 5 | 0.6 | 31 | 3.7 | 36 | 4.3 |
| NUT | 16 | 1.0 | 61 | 4.0 | 77 | 5 |
| XUT | 30 | 1.8 | 54 | 3.3 | 84 | 5.1 |
| SRAT | 21 | 3.9 | 5 | 0.9 | 26 | 4.8 |
| Sum | 388 |  | 559 |  | 974 |  |

**Table S4.** Overlap between transcripts identified by *de novo* analysis and annotated transcripts

|  | Counts |  | Percent of Class |  |
| --- | --- | --- | --- | --- |
|  | *paf1Δ* vs *WT* |  | *paf1Δ* vs *WT* |  |
|  | Decreased | Increased | Decreased | Increased |
| mRNA | 1100 | 359 | 20.32 | 6.15 |
| snoRNA | 1 | 28 | 1.30 | 27.27 |
| CUT | 72 | 34 | 2.38 | 1.41 |
| SUT | 62 | 30 | 4.14 | 1.89 |
| XUT | 77 | 57 | 2.23 | 1.81 |
| SRAT | 11 | 27 | 0.19 | 5.64 |
| NUT | 112 | 101 | 3.08 | 5.18 |
|  | *paf1Δ trf4Δ* vs *trf4Δ* |  | *paf1Δ trf4Δ* vs *trf4Δ* |  |
|  | Decreased | Increased | Decreased | Increased |
| mRNA | 1519 | 844 | 21.47 | 16.24 |
| snoRNA | 17 | 23 | 3.90 | 24.68 |
| CUT | 294 | 70 | 22.19 | 4.11 |
| SUT | 193 | 85 | 19.86 | 6.86 |
| XUT | 372 | 172 | 18.53 | 7.91 |
| SRAT | 45 | 147 | 5.83 | 33.46 |
| NUT | 426 | 229 | 31.72 | 11.27 |

**Table S5.** Counts of transcripts identified by the *de novo* analysis that fall into various categories based on position relative to mRNA coding regions

|  | *paf1Δ* vs *WT* |  | *paf1Δ trf4Δ* vs *trf4Δ* |  |
| --- | --- | --- | --- | --- |
|  | Decreased | Increased | Decreased | Increased |
| sense mRNA Overlap | 872 | 275 | 1111 | 687 |
| antisense mRNA overlap | 23 | 47 | 256 | 353 |
| no overlap with mRNA | 39 | 12 | 151 | 88 |
| Sum | 1268 |  | 2646 |  |

**Table S6.** Counts of antisense transcripts overlapping with various transcript classes.

| Input: Downregulated Antisense Transcripts (n=256) |  |  |  |  |  |  |  |
| --- | --- | --- | --- | --- | --- | --- | --- |
|  | SRAT | CUT | SUT | NUT | XUT | snoRNA | Count |
| cluster 1 | 4 | 14 | 11 | 28 | 32 | 0 | 43 |
| cluster 2 | 11 | 9 | 11 | 36 | 18 | 0 | 86 |
| cluster 3 | 10 | 41 | 36 | 72 | 77 | 0 | 103 |
| cluster 4 | 4 | 8 | 11 | 31 | 24 | 0 | 65 |
| cluster 5 | 4 | 33 | 13 | 37 | 30 | 0 | 50 |
| sum | 33 | 105 | 82 | 204 | 181 | 0 | 347 |

| Input: Upregulated Antisense Transcripts (n=353) |  |  |  |  |  |  |  |
| --- | --- | --- | --- | --- | --- | --- | --- |
|  | SRAT | CUT | SUT | NUT | XUT | snoRNA | Count |
| cluster 1 | 25 | 6 | 2 | 19 | 14 | 0 | 94 |
| cluster 2 | 35 | 10 | 13 | 21 | 33 | 0 | 84 |
| cluster 3 | 20 | 12 | 12 | 31 | 21 | 0 | 95 |
| cluster 4 | 44 | 2 | 9 | 16 | 21 | 0 | 110 |
| cluster 5 | 15 | 8 | 7 | 16 | 36 | 0 | 75 |
| sum | 139 | 38 | 43 | 103 | 125 | 0 | 458 |

